# Mutation bias is a key predictor of adaptation rate

**DOI:** 10.1101/2025.09.22.677663

**Authors:** Shazia Parveen, Aakanksha Madhwal, BK Ruchith, Mrudula Sane, Deepa Agashe

## Abstract

A growing body of work suggests that mutation bias — whereby some types of mutations occur more often than others — can influence the genetic basis of adaptation and the distribution of fitness effects of new mutations. For instance, the loss of specific DNA repair genes shifts the mutation spectrum from a transition-biased to a transversion-biased state in *Escherichia coli*. This reversal of the ancestral mutation bias enhances access to under-sampled variants, increasing the beneficial mutation supply. But are these effects strong enough to alter adaptive outcomes? We show that mutation bias reversal indeed increases adaptation rate in *E. coli* by about 5%, and halves the variation in adaptation rates across replicate populations. Even though replicate populations of a strain diverge and adaptation slows down over time, the effect of beneficial supply on adaptation rate gets stronger. The advantage of bias reversal accrues due to greater beneficial mutation supply, with ∼2-3 times as many beneficial mutations as strains with reinforced bias. Overall, mutation bias strongly influences the genetic basis of adaptation and alters the types of accumulated genomic mutations that may shape subsequent evolution. Our results show that mutation bias is important for predicting evolution, and highlight multiple ways in which mutational processes can shape the genetic and phenotypic outcomes of adaptation.

**SIGNIFICANCE STATEMENT:** Predicting which populations will adapt faster in new environments, and via what kinds of genetic changes, are important questions in evolutionary biology. Recent work suggests that apart from selection, details of the mutational process are also important. Allowing different bacterial strains to evolve in the laboratory, we show that accessing previously poorly sampled mutation types through mutation bias reversal can boost adaptation, leaving clear genomic signatures with at least twice as many beneficial mutations. Moreover, bias favoring specific types of mutations leaves strong imprints in the genome, with different bacterial strains adapting via distinct genetic routes. Knowing the mutation bias thus increases our ability to predict evolution.

## INTRODUCTION

The number of possible mutations that could occur in a genome is vast, but different types of mutations do not occur with equal frequency. For instance, transition mutations — where a purine or pyrimidine base changes to another base of the same type — are more common than transversions, which convert a purine to pyrimidine or vice versa (1–3). If mutation occurrence were unbiased, we would expect about 33% of all single-base substitutions to be transitions. However, this fraction is usually much higher: e.g., in *Escherichia coli,* about 54% of single base mutations are transitions, and for some bacteria it is as high as 97% (4–7). Recent work shows that such biased mutation spectra can have important evolutionary consequences: the underlying mutation bias can determine the genetic basis of adaptation, drive parallel genetic changes across populations, and shape genome evolution across diverse population genetic contexts (8–13). Beyond influencing the genetics of adaptation, mutation bias can also affect the distribution of fitness effects (DFE) of new mutations (14–16). In particular, reversing an existing mutation bias can increase the fraction of beneficial mutations (f_b_) by allowing the population to sample previously rare mutation types (14, 15). Adaptive walk simulations confirmed a general advantage of opposing (i.e., reducing or reversing) the ancestral mutation bias (14). More recent work showed that non-hypermutator cancers acquire driver mutations through mutation bias reversal, further supporting the adaptive benefit of reversing an established bias (17). The increase in the supply of beneficial mutations (S_b_), driven by higher f_b_, implies that reversing an ancestral mutation bias (e.g., from a transition-biased to a transversion-biased spectrum) should facilitate adaptation. In contrast, reinforcing a long-standing bias (e.g., strengthening an existing transition bias) should reduce f_b_, hindering further adaptation (Fig. 1A) (14, 15).

**Figure 1:**
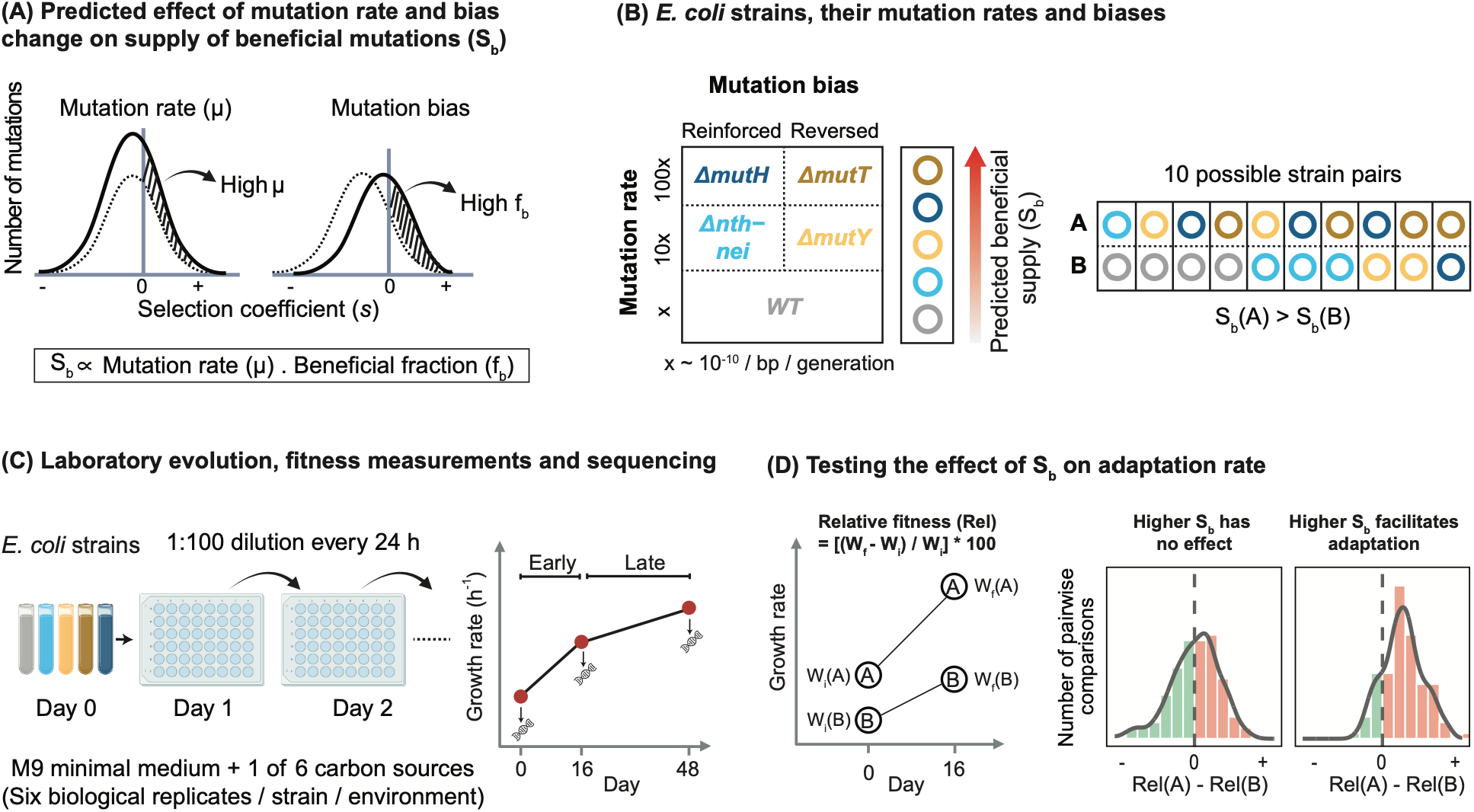
Design of Experiment 1. **(A)** Predicted effects of mutation rate and mutation bias on the supply of beneficial mutations (S_b_). An increase in the overall mutation rate (µ) elevates S_b_ by increasing the total number of mutations generated. In contrast, shifts in mutation bias alter the distribution of fitness effects (DFE), particularly the fraction of beneficial mutations (f_b_). Reversal of an existing mutation bias shifts the DFE rightward, increasing f_b_ and thereby S_b_, whereas reinforcement of the bias shifts the DFE leftward, reducing f_b_ and S_b_. **(B)** *E. coli* strains used in this study, grouped by their mutation rates and mutation biases. WT *E. coli* has a moderate transition bias. Reinforced: more transitions than WT; reversed: more transversions than WT. The schematic shows the predicted rank order of S_b_ (see panel A). Accordingly, for each pair of strains in our study, we predicted the relative adaptation rate. **(C)** To test our predictions, we conducted laboratory evolution in M9 minimal environments supplemented with one of six carbon sources (galactose, gluconate, glucose, glucuronate, N-acetylglucosamine (NAG), or succinate), followed by growth rate measurements and whole genome sequencing. **(D)** Illustration of predicted outcomes of experimental evolution in strain pairs with varying S_b_. We analysed the distribution of pairwise differences in adaptation rates; red bars indicate cases where the strain with higher S_b_ adapted faster (i.e., shows higher relative fitness), and green bars indicate the opposite. The dashed line marks no difference.

To test this prediction, we deleted specific DNA repair genes to generate *E. coli* mutator strains with extreme biases favouring either transitions or transversions (15, 18, 19). Compared to the wild type (WT) (which has a moderate transition bias, with 54% transitions), our mutant strains had biases ranging from 96% transitions to only 2% transitions (15) (Fig. 1B, Table S1). However, deleting DNA repair genes also increases the mutation rate (µ), which independently increases the overall beneficial mutation supply, S_b_ (Fig. 1A) (20–22). Hence, in mutators, the beneficial effects of increased mutation rate and reversed mutation bias are confounded. We therefore focused on pairwise comparisons between strains to test the effect of S_b_ on adaptation rate. Because S_b_ is directly proportional to both f_b_ and µ (S_b_ ∝ µ.f_b_), the relative changes in these quantities should predict the relative advantage to a mutator via either mechanism (Fig. 1A). In our mutators, the fold change in µ is much greater (about 10x and 100x, Table S1) than that in f_b_ (maximum 1.8x in minimal Glucose media, Table S1 (15)), leading to the simple prediction that increasing µ should have a stronger effect on adaptation rate than increasing f_b_ via a bias reversal. Additionally, for a given µ, strains with a reversed bias should have an adaptive advantage compared to strains with reinforced bias. Thus, we expected mutators to gain a large advantage due to high mutation rate (*ΔmutT* and *ΔmutH* outperforming *ΔmutY* and *Δnth-nei* respectively), and an additional advantage due to bias reversal (*ΔmutT* and *ΔmutY* outperforming *ΔmutH* and *Δnth-nei* respectively). Together, these comparisons predict the overall rank order of adaptation rate as: *ΔmutT > ΔmutH > ΔmutY > Δnth-nei* > WT, with a total of 10 possible pairwise strain comparisons (Fig. 1B).

We allowed replicate populations of each strain to evolve in six environments, each containing a single carbon source: galactose, gluconate, glucose, glucuronate, N-acetylglucosamine (NAG), or succinate (Experiment 1, Fig. 1C). We tested our hypothesis by analysing the adaptation rate of each population after 16 days of evolution (early phase, approximately 120 generations) and after 48 days (late phase, about 360 generations), comparing adaptation rates for each strain pair (total 60 pairs across 6 environments). We expected the strain with a higher S_b_ (either due to a mutation rate increase or bias reversal or both) to adapt faster, leading to a right-shifted distribution of differences in adaptation rate (strain with high S_b_ – low S_b_) across all environments (Fig. 1D).

## RESULTS

### Higher supply of beneficial mutations leads to faster adaptation

During early evolution (day 0–16), most populations adapted to the respective carbon source, with an 8%–163% increase in growth rate (Experiment 1, Fig. 2A, Fig. S1A). While growth rate continued to increase during late adaptation (day 16–48), the rate of increase was significantly lower, ranging from 3%–40% (Fig. S1B-C). Using colony counts for a subset of populations, we confirmed that population size also increased during adaptation (Fig. S1D). These results enabled us to test our primary hypothesis: (1) strains with high S_b_ should adapt more rapidly, (2) leading to a right-shifted distribution of pairwise differences in adaptation rates (high S_b_ – low S_b_, Fig 1D). Our experimentally obtained estimates of f_b_ for all strains in glucose media (15) allowed us to directly test point (1). Strains with higher S_b_ values adapted faster in glucose only in the late phase, but not in the early phase (Fig. S2). For the other environments — for which we did not have f_b_ measurements — we used data from glucose (which showed broadly similar patterns to LB media (15)) to derive a qualitative ranking of S_b_ for each strain pair, as described in the Introduction.

**Figure 2.**
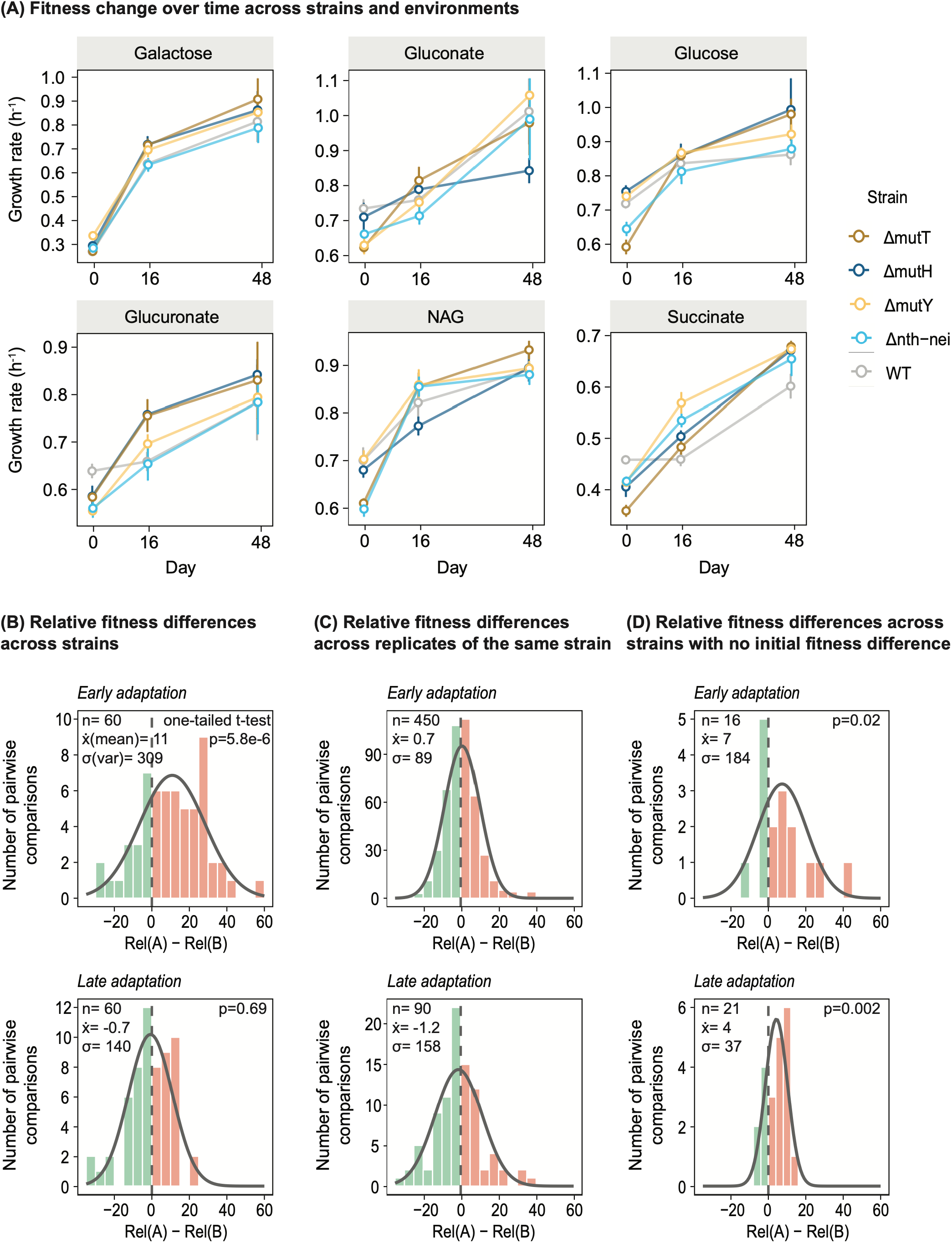
Increased S_b_ facilitates adaptation. **(A)** Observed change in growth rate change over time, measured on day 0, 16 and 48 (Datasheet S1). Points represent mean ± SD (n=6 biological replicates at day 0 and day 16; n=3 at day 48). **(B)** Histogram showing the distribution of differences in adaptation rate (high S_b_ − low S_b_) across all strain pairs during early (days 0–16) and late adaptation (days 16–48). Adaptation rate was calculated as percent fitness increase: ((*final growth rate* − *initial growth rate*)/ (*initial growth rate*)) × 100. Red bars indicate cases where the strain with higher S_b_ adapted faster (i.e., showed higher relative fitness), and green bars indicate the opposite. The dashed line marks no difference; the overlaid curve represents a fitted normal distribution. Sample size (number of pairs, n), mean (x̅), and variance (σ) are given in each plot. A one-sample, one-tailed t-test was used to assess whether each distribution was significantly right-shifted from 0. **(C)** Histogram showing the distribution of adaptation rate differences between all possible pairs of independently evolved biological replicates of a given strain in a given environment, providing a measure of within-strain, across-replicate variability. **(D)** Same as (B), but restricted to strain pairs with no significant difference in initial fitness (determined by Welch’s t-tests, p ≥ 0.05).

Compiling data across all strain pairs and environments, we indeed observed a significant right shift in the distribution of adaptation rate differences during early adaptation (one sample one-tailed t-test, t = 4.79, df = 59, p = 5.78e-06, effect size: Cohen’s d = 0.6) (Fig. 2B). This indicates that strains with a higher S_b_ consistently adapted faster — on average 11% faster — than low-S_b_ strains during early adaptation. During late adaptation, the distribution was not significantly right-shifted (one-sample one-tailed t-test, t = −0.49, df = 59, p = 0.69, Cohen’s d = 0.06). The variance in the distribution of adaptation rate differences during early adaptation was twice as large as that during late adaptation (F-test, F = 2.2058, p = 0.003; compare the early vs. late distributions in Fig. 2B). Thus, strains exhibited both larger and more variable differences in adaptation rate in the early phase. Importantly, the observed S_b_-associated differences in adaptation rate across strains exceeded differences across replicate populations of a given strain during early adaptation (about 3x higher; compare top panels in Figs. 2B and 2C; F-test, F = 3.47, p = 9.99e-14). However, during late phase adaptation, divergence across replicates increased, becoming comparable to between-strain differences (compare bottom panels in Figs. 2B and 2C; F-test, F = 0.89, p = 0.63). Finally, strains with a reversed mutation bias (*ΔmutT* and *ΔmutY*) exhibited 2-3 fold lower variability in adaptation rates across replicate populations compared to strains with reinforced bias (*ΔmutH* and *Δnth-nei*), except for similar variance in *ΔmutT* and *ΔmutH* during late adaptation (Fig. S3). Together, these results suggested that bias reversal repeatedly allowed rapid and parallel adaptation, whereas trajectories were more variable in strains with reinforced mutation bias.

We tested whether differences in adaptation rates could explain differences in initial population size across strains (estimated by counting colony forming units, CFU/mL), which could also generate variation in the mutation supply and affect adaptation rates (23). Most strain pairs (44 of 60) had similar population sizes (Welch’s t-test, p ≥ 0.05; mean ratio ± 95% CI: 1.28 ± 0.23), and adaptation rate differences did not correlate with population size ratios during either early or late adaptation (Spearman’s correlation, early: R = −0.07, p = 0.59; late: R = −0.05, p = 0.72; Fig. S4). Thus, variation in population size cannot account for the observed differences in adaptive outcomes across strains.

However, the effect of S_b_ on adaptation rate described above could be confounded by initial fitness, which influences adaptation rates independently of beneficial mutation supply. Typically, the effect size of beneficial mutations decreases as initial fitness increases, referred to as global epistasis (24–26). Consistent with this pattern, strains with higher initial fitness generally adapted more slowly during both early and late adaptation (significant negative correlations in 5/6 and 4/6 environments in early and late phase respectively; Fig. S5). However, initial fitness did not vary systematically with either mutation rate or mutation bias (Fig. S6), suggesting that the observed association between S_b_ (via mutation bias or rate) and adaptation rate is unlikely to be explained by initial fitness differences. To disentangle the effects of S_b_ from those of initial fitness, we compared adaptation rates for a given strain (i.e., populations with identical mutation rate and bias) across environments to determine the magnitude of global epistasis as a function of initial fitness in different carbon sources. We observed a strong negative correlation between the initial fitness of a strain in a given environment and its early adaptation rate (Fig. S7). However, this pattern was largely driven by a single environment (galactose), in which strains exhibited both the lowest initial fitness and the highest adaptation rate. On excluding galactose, the correlations became weaker and inconsistent across strains (Fig. S7). Similarly, during late phase adaptation, global epistasis was observed overall, but this pattern was driven by succinate populations in *ΔmutT* and *ΔmutH* (Fig. S7). These results suggest that global epistasis did not strongly influence relative adaptation rates. Next, to specifically evaluate the effect of S_b_, we focused on strain pairs with statistically indistinguishable initial fitness. Despite the substantial reduction in sample size, the positive effect of S_b_ on adaptation rate (Fig. 2B) persisted in early adaptation (mean = +7%, one-sample, one-tailed t-test, t = 2.17, df = 15, p = 0.02, Cohen’s d = 0.5), and became significant as well as stronger in the late phase (mean = +4%, one-sample, one-tailed t-test, t = 3.23, df = 20, p = 0.002, Cohen’s d = 0.7; compare Cohen’s d for effect sizes) (Fig. 2D). Thus, even after controlling for differences in initial fitness, high S_b_ increased adaptation rate.

### Strains with a reversed mutation bias adapt faster than those with reinforced bias

In the analysis above, we did not have sufficient statistical power to test the independent effects of mutation bias vs. rate on adaptation. Hence, we performed an additional laboratory evolution experiment in which we evolved 12 distinct clones of *ΔmutT* (reversed bias) and *ΔmutH* (reinforced bias) in glucose minimal medium for 16 days, with five biological replicates per clone (Experiment 2, Fig. 3A-B). These clones were generated by repeated gene knockout experiments using phage transduction, producing strains with similar mutation rates (100x higher than WT) and strain-specific biases, but differing in the number and identity of secondary background mutations that altered initial fitness (Fig. 3A, Datasheet S2). This led to a total of 144 possible pairwise comparisons between *ΔmutT* and *ΔmutH*, allowing us to test the effect of bias reversal with adequate statistical power. As expected, we found a significant right-shift in the distribution of adaptation rate differences between *ΔmutT* and *ΔmutH* clones when considering all possible pairs (one-sample, one-tailed t-test, t = 5.31, df = 143, p = 2.03e-7, Cohen’s d = 0.4) (Fig. 3C). Importantly, this result persisted when considering only the 85 pairs in which clones started with equivalent initial fitness, with an average 5% faster adaptation in *ΔmutT* than *ΔmutH* clones (one-sample, one-tailed t-test, t = 3.78, df = 84, p = 0.0001, Cohen’s d = 0.4) (Fig. 3D). Thus, strains with a reversed bias adapted faster than those with a reinforced bias, after controlling for both mutation rate and initial fitness. As with the previous experiment, the variance of the distribution of adaptation rate differences between strains was significantly greater than that among biological replicates of a given clone (F test, F = 0.46, p = 1.17e-7). Further, consistent with the results from Experiment 1, replicate populations of *ΔmutH* clones had more variable adaptation rates compared to *ΔmutT* replicates (F test, F = 0.49, p = 0.0001, Fig. 3E), indicating access to fewer beneficial mutations. Finally, we measured the effect of global epistasis in our dataset by comparing adaptation rates across clones of the same strain (which have similar mutation rate and mutation bias). Consistent with global epistasis, initial fitness was negatively correlated with adaptation rate for *ΔmutT* clones, but not so for *ΔmutH* clones (Fig. S8). A linear model fit to the complete dataset did not show significant effects of initial fitness, strain, or their interaction on adaptation rate (p>0.29 in each case; Fig. S8), indicating an overall weak effect of global epistasis.

**Figure 3.**
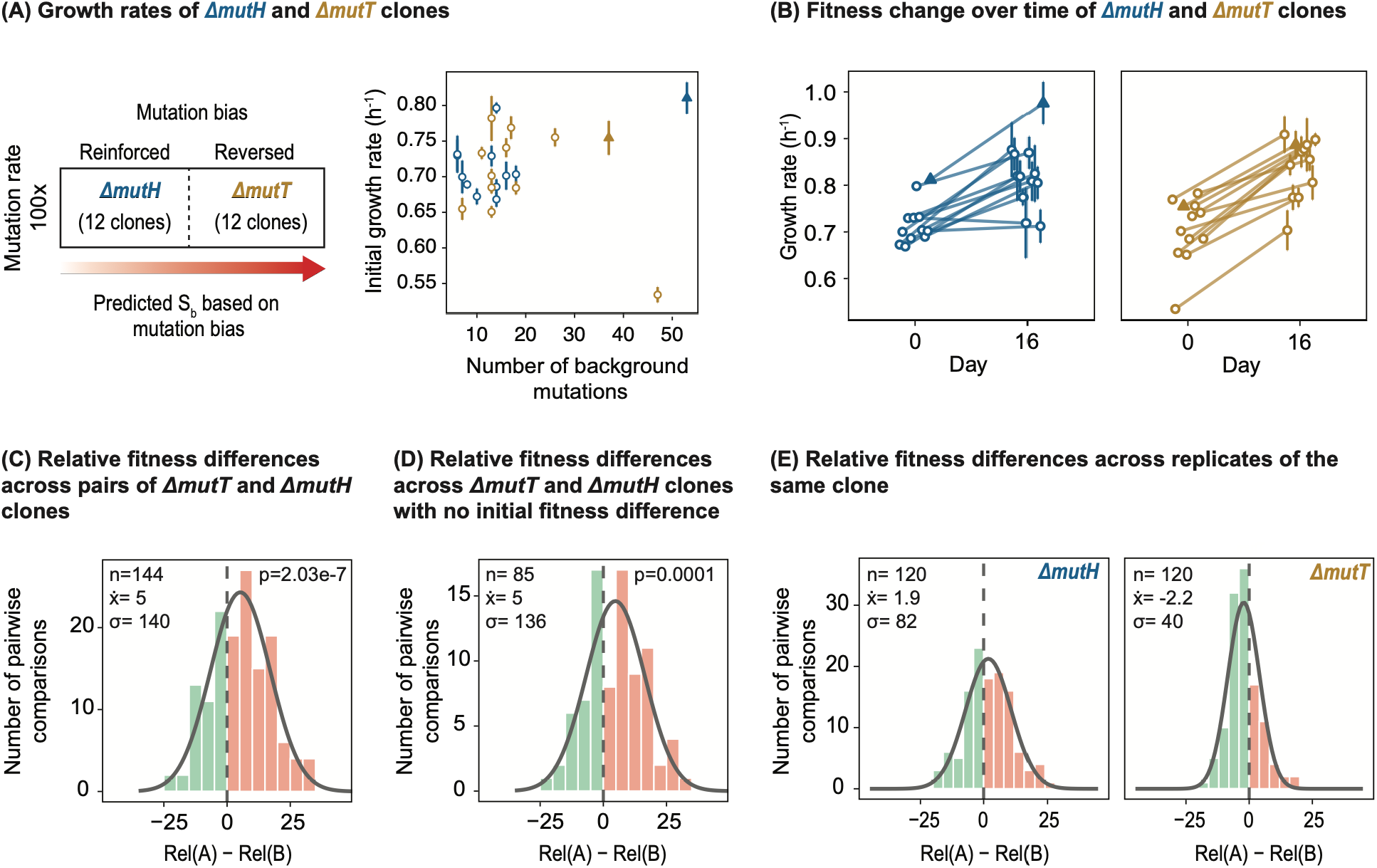
Bias reversal facilitates adaptation (Experiment 2). Twelve independently derived clones each of *ΔmutT* and *ΔmutH* were generated, producing strains with similar mutation rates and biases but differing in the number and identity of background mutations. **(A)** Initial fitness (growth rate) of mutator clones, measured in M9 minimal medium with glucose (mean ± SE, n=3-6 technical replicates per clone), as a function of the number of background mutations in each clone. Spearman’s rank correlation (*ΔmutT*: R = 0.019, p = 0.96; *ΔmutH*: R = −0.2, p = 0.56). Clones that were also used for Experiment 1 (results in Fig. 2) are indicated by filled triangles. **(B)** Change in the growth rate of each clone after 16 days of evolution in M9 minimal medium with glucose (n=5 independently evolved biological replicates per clone). Each point represents mean ± SD for a biological replicate (n=3 technical replicates). **(C)** Histogram showing the distribution of differences in adaptation rates between each pair of *ΔmutT* (reversed bias) and *ΔmutH* (reinforced bias) clones. **(D)** Same as (C), but restricted to clone pairs with no significant difference in initial fitness. **(E)** Histograms showing the distribution of growth rate differences between all possible pairs of biological replicates of the same clone of each strain, providing a measure of within-clone across-replicate variability. In panels **C-E**, red bars indicate cases where the strain with higher S_b_ adapts faster (i.e., shows higher relative fitness), and green bars indicate the opposite. The dashed line marks no difference; the overlaid curve represents the fitted normal distribution. Sample size (number of clone pairs, n), mean (x̅), and variance (σ) are indicated within each plot. For panels C and D, one-sample one-tailed t-tests were used to assess whether each distribution was significantly shifted to the right relative to 0.

### Strains with reversed bias sample more beneficial mutations

To investigate genomic signatures associated with adaptation, we sequenced day 16 and day 48 evolved populations from Experiment 1, including all replicates of WT, *Δnth-nei*, *ΔmutY*, *ΔmutH*, and *ΔmutT* strains propagated across each of the six environments (Fig. 1C, Datasheet S3). We compared each population with its respective ancestral population to identify *de novo* mutations acquired during adaptation and test the effect of predicted differences in S_b_ on accumulated mutations.

We first analysed the total number of mutations observed in each population (Fig. 4A), which is a function of the mutation rate as well as selection and linkage between mutations with different selection coefficients. Additionally, new mutations in DNA repair genes — acquired during adaptation — could alter the mutation rate and influence the total number of mutations. We identified 6 out of 360 populations with significantly more mutations (outliers beyond >1.5x IQR for a given strain and environment), though we did not find mutations in known DNA repair genes in all of them. The outliers included two NAG-evolved *Δnth-nei* populations, two succinate-evolved *ΔmutY* populations, one succinate-evolved WT population, and one gluconate-evolved *ΔmutY* population, all at day 16 (Fig. 4A). The small number of outliers, together with a lack of an environment- or strain-specific pattern of outlier occurrence, suggested that mutation rate was largely stable and mutation rate changes did not systematically alter experimental outcomes. At day 16, the total number of mutations increased with mutation rate (Fig. S9, Table S2). All mutators had more mutations than WT (Table S3), but there was no difference between strains with 10x vs. 100x higher mutation rates than WT (Table S3). Similarly, mutation rate did not explain the number of mutations at day 48 (Fig. S9, Table S2). Instead, at day 48, mutation counts were consistently higher for reversed bias strains for a given mutation rate (Fig. 4A). Thus, the total number of mutations was governed primarily by selection and linkage between mutations, rather than mutation rate.

**Figure 4.**
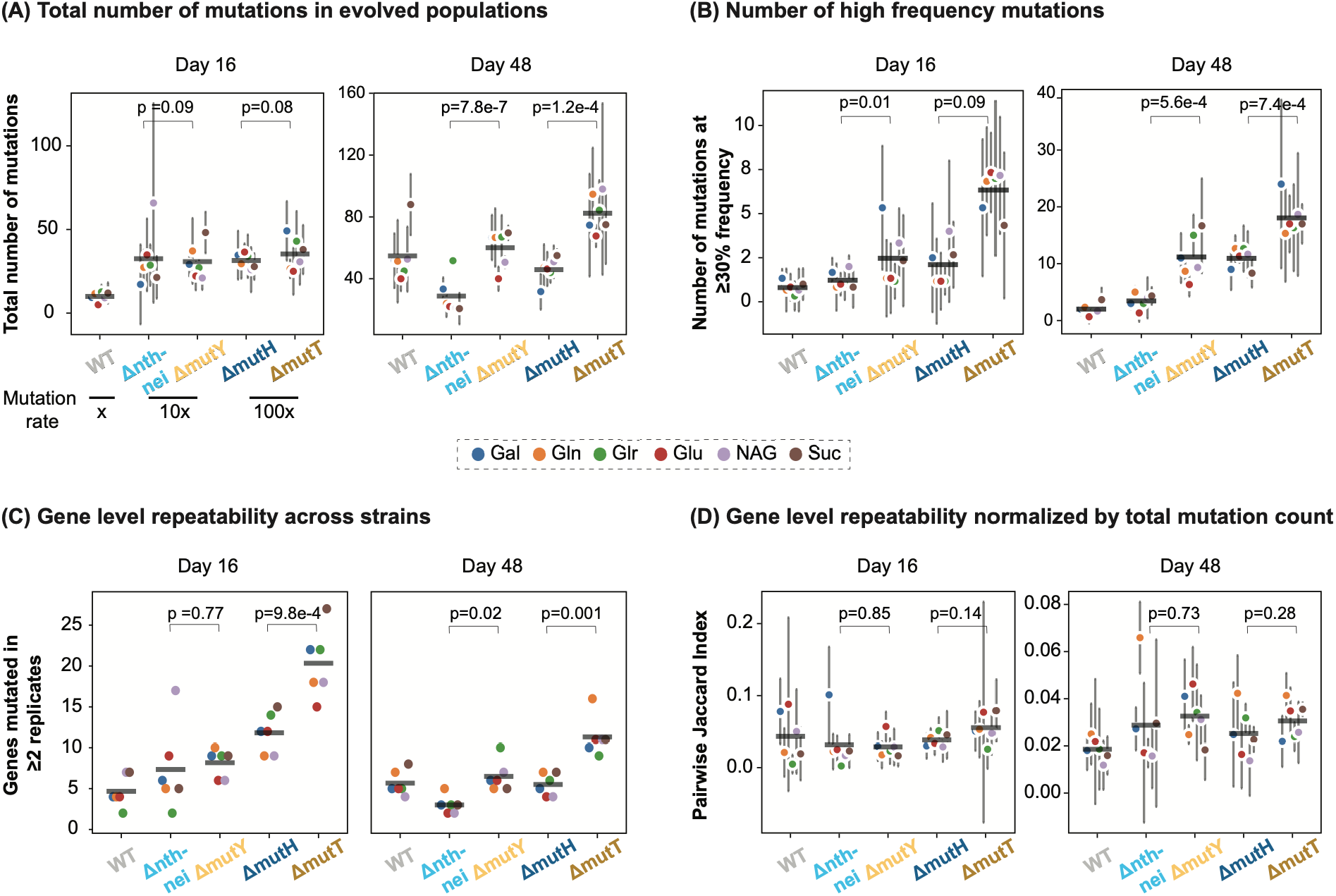
Strains with reversed mutation bias sample more beneficial mutations. **(A)** Total number of mutations found in each strain across different environments, shown as mean ± SD across replicate populations within each environment. Different colours represent different environments. **(B)** Number of high-frequency mutations (≥30%), mean ± SD for a given strain across replicate populations within each environment. **(C)** The number of genes that acquired mutations in at least two independent replicate populations of the same strain evolving in a given environment. **(D)** Pairwise Jaccard similarity index calculated for each strain within each environment, defined as the number of shared genes divided by the total number of unique genes mutated across two populations. Each point represents the mean ± SD for a given strain across replicate pairs within each environment. At day 16, six replicate populations per strain yield 15 pairwise comparisons, whereas at day 48, three replicates yield three pairwise comparisons. In panels A and B, p values were derived from fitting GLMs with negative binomial errors: Number of mutations ∼ Strain * Environment (see full model output in Datasheet S4 and S5). In panels C and D, p values were derived from paired t-tests.

As expected, most mutations were rare, and few mutations (<18% of total) reached frequencies ≥30% (Fig. S10). Adaptive mutations are more likely to reach high frequencies, so we expected strains with higher S_b_ to have a greater number of such mutations. However, some neutral or mildly deleterious mutations can also rise in frequency by hitchhiking with adaptive mutations — an effect that is expected to increase with mutation rate. Therefore, we focused on comparisons across strains with similar mutation rates, allowing us to test the effect of mutation bias shifts. Strains with reversed bias consistently accumulated 2-3 times more high-frequency mutations (≥30% frequency in a population) than those with reinforced bias (Fig. 4B), consistent with the predicted increase in S_b_ due to bias reversal. Interestingly, *ΔmutY* (µ = 10x, reversed bias) had as many high frequency mutations as *ΔmutH* (µ = 100x, reinforced bias) on both day 16 and 48 (Fig. 4B). Thus, the effect of shifting the distribution of fitness effects via bias reversals could be as large as the effect of a 10-fold increase in mutation rate (27).

Another hallmark of highly beneficial mutations is their tendency to be selected repeatedly in independently evolving populations (28, 29). Most mutations in our dataset were unique to individual populations, but approximately 6% of the 8,527 (i.e., about 506) genes carrying new mutations were mutated in at least two replicate populations. This gene level repeatability was two-fold higher in strains with reversed mutation bias by day 48 (Fig. 4C), with similar patterns for mutation level repeatability (about 335 mutations were shared across replicates, <4% of all mutations; Fig. S11A). These results were consistent with the idea that bias reversal increases the supply of beneficial mutations. However, higher repeatability could also reflect differences in the total number of mutations across strains: the higher the total number of mutations, the higher the probability of finding shared mutations across replicates. Hence, we calculated pairwise Jaccard similarity between populations, which accounts for total mutation counts. Consistent with increasing genetic divergence over time (30), the Jaccard index decreased significantly from day 16 to 48 at both gene level (paired t-test: t = 2.613, df = 29, p = 0.014; note the distinct y-axes, Fig. 4D) and mutation level (paired t-test: t = 3.173, df = 29, p = 0.003, Fig. S11B) (30). However, the index had comparable values across strains in both cases (Fig. 4D, Fig. S11B), suggesting that the increased parallelism in bias-reversed strains arose primarily due to access to a larger pool of adaptive mutations, rather than changes in proportional overlap in genes or mutations.

Finally, we estimated the distribution of selection coefficients (s) for each evolved population by analysing the change in allele frequencies (mutation frequency) (p) over time (t), using the logistic model: *dp*/*dt* = *sp*(1 − *p*). The model assumes an infinitely large population, no genetic drift, no linkage, and constant selection. Some of these assumptions (e.g., no linkage) do not hold in our populations. Nonetheless, this analysis allows a coarse-grained estimation and comparison of DFEs across strains, which is instructive. The majority of mutations detected at day 16 had negative selection coefficients and were purged by day 48, consistent with strong purifying selection (Fig. S12A). Notably, strains with reversed mutation bias had a higher fraction of mutations with positive selection coefficients (*ΔmutY*: 12%, *ΔmutT*: 16%) compared to strains with reinforced bias (*Δnth-nei*: 4%, *ΔmutH*: 13%) (Fig. S12B), supporting our inference that bias reversal confers an adaptive advantage.

### Mutation bias determines the spectrum of mutations accumulated during adaptation

Several retrospective analyses show that mutation bias can strongly shape the genetic basis of adaptation (8, 9, 31, 32). Here, we directly tested this idea for several strains and environments. The overall mutation biases (i.e., transition-transversion bias, single nucleotide polymorphism (SNP)-indel bias, GC to TA bias, non-synonymous bias and coding bias) and mutation spectrum of single base changes in evolved populations strongly reflected the baseline spectrum of the respective strains under genetic drift (Fig. S13A-F; see supplementary materials and methods for corrected estimates of mutation bias following (15)). Focusing on the transition-transversion bias, we found that high-frequency mutations (≥ 30%) in all populations were highly enriched in the mutation type favoured by the underlying bias (Fig. 5A). Thus, mutation bias leaves strong genomic signatures even under short-term rapid selection, consistent with previously observed patterns across thousands of generations of evolution (31).

**Figure 5:**
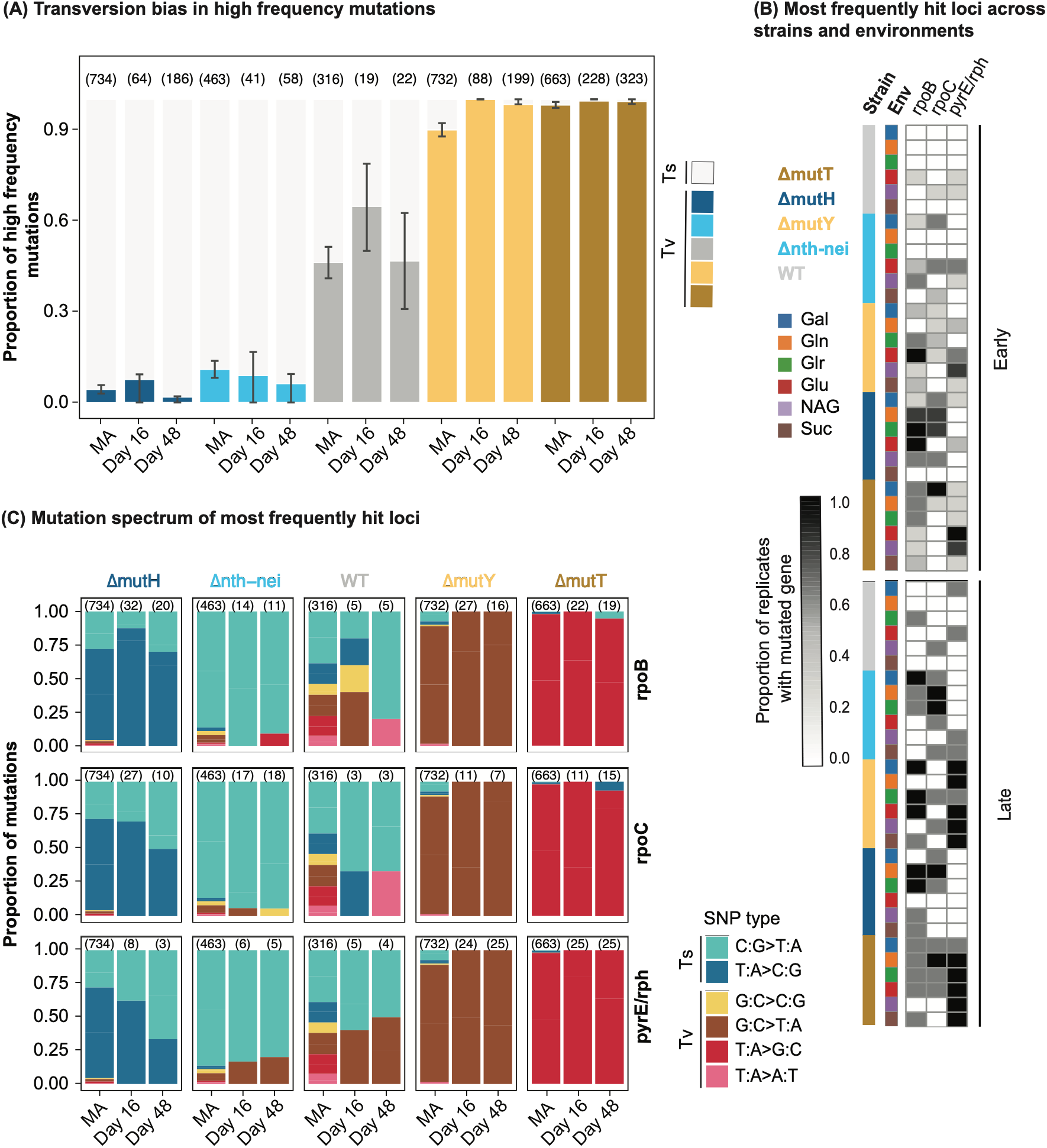
Mutation spectrum of putatively adaptive mutations. **(A)** Transversion bias (Tv bias) in high-frequency mutations. For each strain, the leftmost bar shows the Tv bias under genetic drift (under mutation accumulation, MA; data from (15), but see supplementary materials and methods for corrected estimates of mutation bias), followed by data for day 16 and day 48 of evolution. For MA, error bars represent 95% confidence intervals; for evolved lines, error bars indicate the IQR of Tv bias across different environments. Numbers in parentheses represent the number of point mutations used to infer Tv bias. **(B)** Most frequently mutated loci (*rpoB, rpoC, pyrE/rph*) during experimental evolution. The grayscale intensity indicates the proportion of replicates in which a given gene was mutated. (**C)** The spectrum of mutations observed at the loci shown in panel B, compared to the mutation spectrum observed under MA. Cool colours indicate transitions; warm colours indicate transversions.

Next, we focused on the most frequently mutated loci in our dataset — *rpoB* and *rpoC*, and the intergenic region of *pyrE/rph* — which acquired mutations in multiple strains across different environments (Fig. 5B, see Fig. S14 for the full heatmap). These loci are often mutated in laboratory evolution experiments and are broadly advantageous (33). For example, mutations at 12 of 49 unique amino acid positions in *rpoB* and 2 of 44 unique mutated positions in *rpo*C in our dataset were involved in adaptation in prior studies (34). A deep mutational scanning study of *rpoB* included 11 of our observed amino acid substitutions, and all but one were beneficial in several minimal media (35) (Fig. S15). At the *pyrE/rph* intergenic region, some of the observed mutations likely compensate the *rph* frameshift mutation (found in all our strains) that was inherited from the *E. coli* K-12 MG1655 ancestor, which affects transcription-translation coupling in the *pyrE/rph* operon (36) (Fig. S16). In our study, mutations at all three frequently hit loci rose to average allele frequencies of about 27% on day 16 and 42% on day 48, with most mutations in *rpoB* (168 of 173) and *rpoC* (122 of 123) being nonsynonymous (Fig. S17A). Thus, these mutations are likely beneficial, allowing us to analyse the effect of underlying mutation bias on the genetic basis of adaptation. While mutations at each locus occurred at similar positions across strains (Fig. S17B), the specific base pair substitutions observed in evolved lines strongly matched the baseline mutation spectrum of each strain (Fig. 5C). Further, each strain was enriched for distinct types of amino acid substitutions, with potential implications for protein function and subsequent evolution. For instance, acidic-to-basic substitutions in *rpoB* were enriched in *ΔmutH* and *Δnth-ne*i but absent in *ΔmutT* and *ΔmutY*, whereas polar-to-basic substitutions in *rpoC* were enriched in *ΔmutT* and *ΔmutY* but not in *ΔmutH* and *Δnth-nei* (Fig. S18). Thus, ‘generalist’ mutations at loci that offer broad fitness advantages in *E. coli* show strong imprints of strain-specific mutation biases. This effect of underlying mutation bias was consistent even for ‘specialized’ adaptive mutations observed in environment-specific targets in galactose and gluconate (Fig. S19).

The distinct types of amino acid substitutions observed in different strains suggests the potential for mutation bias to differentially alter protein function. Even if many mutations in the above analyses were neutral or weakly deleterious in the current experimental regime, they may substantially alter the course of subsequent evolution by determining the nature and frequency of allelic variation present in populations, and the epistatic fitness landscape encountered by new mutations (37). Further, *pyrE/rph* intergenic mutations occurred over 4 times more often in *ΔmutT* and *ΔmutY* than in other strains (Fig. 5C), indicating potentially distinct genetic paths to increased fitness. Thus, even if the impact of mutation bias on adaptation rate is transient, it can dramatically alter the type and nature of mutations segregating in populations, potentially influencing the course of subsequent evolution.

## DISCUSSION

We provide one of the first systematic experimental tests of the role of mutation bias in adaptive evolution, and find that it substantially affects both the rate of adaptation and the genetic changes associated with it. Mutation bias can thus shape how populations traverse fitness landscapes and mutational space, perhaps reaching distinct fitness peaks. As expected from prior work, adaptation rate in our experiments slowed over time (38), with a concurrent reduction in differences between strains and increasing divergence across replicate populations of the same strains (39). Together, these results indicate rapid divergence in adaptation rates and an accompanying increase in evolutionary unpredictability with time. In contrast, the effect of S_b_ became more pronounced over time, suggesting a consistent effect of mutation rate and bias on short-term adaptive outcomes. Additionally, we observed higher variation across replicates of strains with reinforced bias as compared to strains with reversed bias in both experiments (Fig. S3, Fig. 3E), consistent with S_b_ differences: access to fewer beneficial mutations limits overall adaptation and increases variability among replicates, presumably because some replicates do not fix the most advantageous mutations.

By quantifying the effect of mutation bias while controlling for mutation rate and initial fitness, our study also enhances current understanding of the evolution and fate of mutators, which often have altered mutation spectra in addition to higher mutation rates (40). Theory as well as empirical data indicate that mutators should have a broad selective advantage, explaining their recurrent emergence and persistence in natural, laboratory, and clinical microbial populations (41–43). Our work provides experimental support for the theoretical prediction that some mutators may gain an additional advantage due to a reversal of the ancestral mutation bias (44). This advantage could allow such mutators to persist longer than expected, e.g., as observed in some long-term experimental evolution (LTEE) populations (45). We also found that mutators can rapidly acquire secondary (background) genomic mutations, changing initial fitness and thereby influencing adaptation rate via global epistasis, independent of beneficial supply. Thus, our work identifies three distinct mechanisms that govern the fate of mutators and the predictability of mutator evolution: mutation rate (via S_b_), mutation bias (via f_b_ and therefore S_b_), and rapid acquisition of secondary mutations (via initial fitness).

Although direct quantitative comparisons were limited, we find that the effect of global (diminishing returns) epistasis on adaptation rate was generally weak and variable, contrasting the consistent positive effects of mutation rate and bias. Although adaptation rate reduced with higher initial fitness (as expected due to global epistasis), this relationship was largely driven by differences across strains that were confounded with the effect of S_b_, or was influenced by populations adapting to specific environments. Thus, in our experiments, adaptation rate differences between populations were governed more strongly by S_b_ — which determines the rate with which beneficial mutations are sampled — rather than initial fitness, which largely affects the effect size of beneficial mutations (22, 24–26). However, global epistasis likely did contribute to the temporal slowing down of adaptation over time, as observed in prior work (24–26, 38, 46).

Slower adaptation in the later phase of our experiment could also arise due to genetic draft, i.e., deleterious mutations hitchhiking with adaptive alleles (47). Accumulated deleterious mutations can reduce the fixation rate of beneficial mutations, causing adaptation rate to stall and potentially even drive fitness declines (48, 49). Although we did not observe significant fitness declines in our populations, weakly deleterious mutations segregating in our mutator populations may eventually drag them towards adaptation rates of zero well before the beneficial mutation supply is exhausted (50, 51). The number of hitchhiking deleterious mutations and their effects on population fitness trajectories are sensitive to several parameters including the shape of the DFE; and as such, remain difficult to estimate for our populations. Apart from the deleterious load, clonal interference — competition between beneficial mutations arising on distinct genetic backgrounds — can also slow down adaptation rates in large populations, and is likely to play a role in our experiment. Understanding the effects of deleterious load and clonal interference in rapidly adapting mutator populations remains an important avenue for future research.

In summary, our work shows that mutation biases leave strong genomic signatures that can have significant implications for later evolution, e.g., via altered epistasis and the accumulation of distinct types of amino acid substitutions. Therefore, mutation bias may alter the course of subsequent adaptation by influencing the nature and consequences of new mutations. As demonstrated by prior analysis of LTEE lines of *E. coli*, mutators with altered mutation bias can have lasting effects on mutation spectra even under strong selection (31). Our work shows clear evidence for a substantial role of mutation bias in shaping the outcomes of adaptive evolution, complementing growing understanding of the evolutionary role of mutation biases (13, 32, 52). We highlight the need to quantify the effects of mutation bias in diverse contexts across different timescales and organisms, and its consequences for evolutionary dynamics.

## METHODS

For Experiment 1 (Fig. 1B-C), we used the same bacterial strains described in (15). Briefly, *E. coli* K12-MG1655 from the Coli Genetic Stock Centre (CGSC) was streaked on LB (Luria Bertani) agar plates and a random colony was chosen as the WT ancestor (WT) for all experiments. We used mutator strains (*ΔmutT, ΔmutH, ΔmutY, Δnth* and *Δnei*) from the Keio collection (BW25113 background) to generate gene knockouts in our WT ancestor using P1 phage transduction, followed by removal of the kanamycin resistance marker using transformation. We generated the *Δnth-nei* double knockout by first deleting *nth* and then deleting *nei*. Because these whole-gene knockouts immediately increased the mutation rate, each strain accumulated some spontaneous secondary mutations (in addition to the focal DNA repair gene deletion) during kanamycin marker removal (see Datasheet S2 for details of secondary mutations). For Experiment 2 (Fig. 3A), we generated 11 additional clones each of *ΔmutT* and *ΔmutH*, using independent transductant colonies following P1 phage transduction and curing each clone independently, such that each endpoint clone accumulated a unique set of spontaneous secondary mutations during the curing process. We confirmed all strains by PCR (primer details are provided in Table S4).

In Experiment 1 (Fig. 1B-C), we performed laboratory evolution of all strains with 6 biological replicate populations in each of six M9 minimal media environments supplemented with a single carbon source: 5 mM galactose, succinate, glucuronate, gluconate, N-acetylglucosamine, or glucose over 16 days with 1:100 dilution every 24 h, in 48-well microwell plates (200 rpm shaking at 37°C) (see supplementary materials and methods for details). Three randomly chosen replicates of each strain were further evolved until day 48 in each environment, leading to approximately 234, 355, 429, 441, 443, and 286 generations in galactose, gluconate, glucuronate, glucose, NAG, and succinate respectively. Following the transfers, evolving populations were periodically frozen at –80°C; stocks were revived overnight in the respective selection medium before further measurements. We measured exponential growth rates of populations at days 16 and 48, along with their respective ancestral populations, with three technical replicates per population using a Biotek Multimode plate reader (425 cpm, 3 mm, orbital shaking at 37°C). We estimated the growth rate using the Curve Fitter software (53). We confirmed that growth rate measurements across different machines (Spearman’s correlation: R=0.98, p = 4.3e-11) and days (Spearman’s correlation: R=0.89, p = 7.6e-29) were highly repeatable (Fig. S20A-C).

For Experiment 2 (Fig. 3A), we allowed 12 independently derived clones of *ΔmutT and ΔmutH* to evolve in M9 minimal media with 5 mM glucose for 16 days as described above, with 5 biological replicate populations per clone (i.e., total 60 independently evolved populations per strain).

We performed whole-genome sequencing of ancestral and evolved populations of Experiment 1 by reviving them in Luria broth (LB) for 4-5 h and extracting genomic DNA (Qiagen DNeasy Blood & Tissue kit). After checking DNA quality, we prepared libraries using Illumina (M) Tagmentation kit (CAT No: 20060059) and sequenced them on the NOVAseq 6000 platform (paired-end sequencing, 2×100 bp). We used fastp (version 0.24.0) for adapter trimming (54). The sequencing generated high-quality reads with an average depth of 139x per sample (range 70x–255x) and >99% read mapping. Mutation calling was performed using Breseq (version 0.39.0) (55) with default polymorphism parameters (mutations with <5% frequency were discarded as likely false positives). Mutations that were found in the corresponding ancestral populations were filtered out to retain only *de novo* variants in the evolved populations. All downstream analyses were performed using this filtered list of mutations. To estimate the false negative rate of sequencing, we checked calling rates for known mutations: (1) the deleted repair gene in each ancestor (it was called correctly as missing coverage in 100% of samples) and (2) two ancestral mutations in the original WT strain: G→A at position 2845011 (found in all samples at 100% frequency) and a GC insertion at 4296380 (found in all samples at 100% frequency, except for one where it was observed at 88.2%).

## ACKNOWLEDGEMENTS

We thank Lindi Wahl, members of the Agashe lab, and two anonymous reviewers for discussion and comments on the manuscript, Swastik Padhy and Adrita Chakraborty for laboratory assistance, and the NCBS NGS facility for sequencing. We thank Dr. Alaksh Choudhury for sharing the raw fitness data of *rpoB* mutations used in Figure S15.

## Author contributions

DA and MS conceived the project; all authors contributed to experimental design; SP, AM, RBK and MS conducted experiments; SP and AM analysed data; SP prepared figures; DA and SP wrote the manuscript with input from all authors; DA acquired funding.

## Data availability

All raw data underlying figures are available as supplementary files.

## SUPPLEMENTARY MATERIALS

Materials and Methods

Figures S1 to S20

Tables S1 to S4

