## Supplementary information for "Mutation bias is a key predictor of adaptation rate"

### SUPPLEMENTARY MATERIALS AND METHODS

#### Generating experimental strains

We obtained the mutator strains (*ΔmutT, ΔmutH, ΔmutY, Δnth* and *Δnei)* in an *E. coli* K-12 BW25113 background from the Keio collection at Coli Genetic Stock Centre (Yale University) (1). These knockouts were constructed by replacing the open reading frame of the relevant gene with a kanamycin resistance cassette, such that removing the cassette generates an in-frame deletion of the gene. We moved the knockout locus from the BW25113 background to that of our WT i.e., MG1655 using P1 phage transduction (2). Successful transduction would replace the focal DNA repair gene with a kanamycin resistance cassette, resulting in kanamycin resistant colonies on selective Luria agar plates with 50 µg/mL kanamycin. After PCR confirmation using DNA repair gene specific primers (Table S4), positive transductants were grown in luria broth (LB) for 4-5 hours and then frozen with glycerol at -80°C. The kanamycin resistance cassette was cured by transforming the transductants with the pCP20 plasmid that carries the flippase - recombinase enzyme encoding machinery, a temperature sensitive *ori*, and an $Amp^R$ marker. The kanamycin resistance cassette is flanked by flippase recognition sites which act as targets for the enzymes encoded by the plasmid. After incubation with pCP20 at 30°C for 48 hrs, the plasmid was cured by incubating the strains at 37°C for another 48 hrs. Removal of the kanamycin resistance cassette and pCP20 was confirmed by replica plating the strains on both Luria agar plates containing 100 µg/mL carbenicillin (to select for $Amp^R$) and 50 µg/mL kanamycin. Colonies that did not grow in either antibiotic were confirmed using PCR, grown for 4-5 hrs in LB, and saved as glycerol stocks at -80°C.

#### Laboratory evolution

**Experiment 1**: We carried out evolution of all four mutators (*ΔmutT, ΔmutH, ΔmutY, Δnth-nei*) and WT (same bacterial strains as (3)), for 6 biological replicate populations per strain in six M9 minimal media environments, each supplemented with a single carbon source (5 mM galactose, succinate, glucuronate, gluconate, N-acetyl-D-glucosamine, or glucose; Sigma-Aldrich). To obtain starting cultures for experimental evolution, we revived strains by inoculating 2 µL of the frozen glycerol stock in 600 µL LB and incubated for 14 h at 37 °C, with orbital shaking at 200 rpm. This was done in 6 different wells of a sterile 48-well culture plate (Biofil) to get 6 biological replicates. To start experimental evolution, we prepared M9 minimal salts (Difco) containing the respective carbon source. The composition of M9 minimal medium was as follows: for 50 mL of M9 minimal media: 44 mL water, 5 mL of 10X M9 minimal salts

(Difco), 500 µL of 0.5 M carbon source solution (made in water), and 500 µL of 0.4 M $MgSO_4$ solution (Fisher Scientific). We dispensed 600 µL of this medium into each well of a 48-well sterile culture plate. From the revived cultures of mutators and WT, we inoculated 6 µL into alternate wells of the 48-well plate and also made ancestral stocks by mixing 400 µL of culture with 400 µL of 60% glycerol stored at –80°C. The remaining wells were left blank to detect contamination. We used a checkerboard pattern of inoculation to minimize cross-contamination across independently evolving populations. After inoculation, we covered the plate edges with parafilm to prevent contamination and incubated plates at 37 °C for 24 h with orbital shaking at 200 rpm. Every 24 h, we transferred 6 µL of each population to fresh media, maintaining a 1:100 dilution. We carried out daily transfers in this way for 16 days for all six replicates per strain, and continued three randomly chosen replicates (replicates 1, 2 and 6) for 32 more days, till day 48. We prepared glycerol stocks of evolving populations every alternate day, as described for ancestral stocks above.

After 16 days of experimental evolution, we tested for contamination by randomly choosing 1-2 populations of each strain evolved in a given environment, and streaking it on LB and MacConkey (Difco) agar plates to screen for the expected colony morphology typical of our *E. coli* strains. We also confirmed strain identity at day 16, by carrying out PCR on a small aliquot of the frozen population stock for 2-3 populations of each strain in a given environment to detect the expected deletion of the strain-specific focal DNA repair gene. Similarly, after 48 days of evolution, we PCR-tested one biological replicate of each strain in a given environment. We detected cross-contamination in populations of *ΔmutY* and *Δnth-nei* evolving in NAG and gluconate at day 48, and traced it close to the 16-day mark. Hence, we discarded the contaminated stocks and repeated experimental evolution for these populations from day 16 to 48.

**Experiment 2**: We evolved all 12 independently derived clones of *ΔmutT* and *ΔmutH* in M9 minimal medium supplemented with 5 mM glucose for 16 days: one clone per strain from Experiment 1, and 11 additional clones. However, the clones from Experiment 1 had acquired some additional background mutations before we initiated Experiment 2 during the intervening rounds of culturing (see Datasheet S2 for the list of background mutations). Each clone was evolved as five independent biological replicate populations, yielding a total of 60 independently evolved populations per strain, using the same protocol as in Experiment 1. To check for contamination, we screened 19 randomly selected populations from each strain at the end of the evolution experiment (day 16).

**Growth rate measurements**

To measure evolved and ancestral growth rates, we inoculated 6 µL of each population from its glycerol stock into 600 µL of the respective selection medium for 14-16 h, at 37 °C, with orbital shaking at 200 rpm in 48-well plates to get a primary culture. Then, we inoculated 6 µL of this revived culture into 600 µL of the same medium in 48-well plates in triplicates and incubated the plate in a Biotek Multimode plate reader at 37 °C with orbital shaking at 425 cpm, 3mm for 20-24 h. Every 15 minutes, the plate reader measured the optical density (OD) at 600 nm for all wells. Each plate included a reference strain (the parent WT strain) to ensure consistency across plate reader runs, as well as blank control wells to verify media sterility. We estimated the exponential growth rate using the Curve Fitter software (4).

**Estimating the number of generations of evolution**

Since growth rates vary across carbon sources, our strains could have evolved for different numbers of generations in each selective medium, during 48 days of evolution. We therefore estimated the number of generations of evolution in the WT, using the number of doublings per day in the WT ancestor, in each medium. We first measured the number of cells used to found each population. For this, we inoculated 2 µL of frozen stock of the WT strain in 600 µL LB broth in a 48 well microplate. We incubated the plate for 15 h at 37 °C with orbital shaking at 200 rpm. To estimate the number of colony forming units (CFU; cells), we prepared several dilutions of the revived culture in sterile 0.9% NaCl (saline) solution. We took 100 µL aliquots from $10^4$-fold or $10^5$-fold dilutions and spread-plated them on Luria agar plates. We calculated CFUs/mL as: $(number\ of\ colonies\ \times\ dilution\ factor)\ /\ volume\ plated\ (mL)$**.** From this, the total number of CFUs present in a 6 µL inoculum volume was determined, as the starting population size. To estimate the number of cells at the end of the first growth cycle (generation 0), we inoculated 6 µL of the revived WT culture (above) into 600 µL of M9 minimal medium supplemented with 5 mM of each of the six carbon sources in 48-well plates, incubated as above. After 24 h, final CFU/ml was estimated by spread plating 100 µL aliquots from $10^4$-fold or $10^5$-fold dilutions onto Luria agar plates. The number of doublings over the 24 h growth period was calculated using the formula: $Number\ of\ doublings\ =\ log_2\ (Final\ CFU\ /\ Initial\ CFU)$, averaging across three technical replicates. We used the number of doublings per 24 h to estimate the total number of generations of experimental evolution in each medium, by multiplying with the total number of days of evolution (16 or 48).

**Estimating change in population size**

We quantified changes in population size (CFU/mL) over the course of experimental evolution, comparing ancestral (day 0) and evolved (day 48) populations of one biological replicate (with the highest overall adaptation rate) of each strain in two environments: M9 minimal media with galactose or gluconate. For each replicate, frozen stocks from day 0 and day 48 were revived by inoculating 2 µL into 600 µL of M9 minimal medium supplemented with 5 mM of the respective carbon source. Revivals were conducted in 48-well plates and incubated for 15 h at 37 °C with orbital shaking at 200 rpm. 6 µL of the revived culture was transferred into 600 µL of fresh media and grown for an additional 24 h under the same conditions. Serial dilutions of grown cultures were prepared in sterile 0.9% NaCl solution. For CFU counts, 100 µL of either the $10^4$-fold or $10^5$-fold diluted cultures was spread-plated onto M9 agar plates supplemented with the respective carbon source. Each dilution was plated in triplicates and plates were incubated at 37 °C for 18–20 h, after which colonies were counted. Mean CFU/mL across three technical replicates was calculated as described above.

**Whole genome sequencing**

To perform whole genome sequencing, we inoculated 20 µL of each ancestor, 16-day, and 48-day evolved population into 600 µL of LB in 48-well plates. Plates were incubated at 37 °C with orbital shaking at 200 rpm for 4-5 hours. Genomic DNA was extracted from grown cultures using the Qiagen DNeasy Blood & Tissue Kit, quantified using Qubit for ~10 randomly chosen samples for each set of 96 samples that were processed together. We prepared paired-end libraries for each population using the Illumina DNA Prep (M) Tagmentation kit (96 Samples, CAT No: 20060059). Sequencing was performed on the NovaSeq 6000 platform using 2x100 paired-end reads.

**Data analysis**

**Correcting mutation rate and mutation bias estimates using data from Sane et al., (2025)**

Sane et al., 2025 (3) performed mutation-accumulation (MA) experiments using the same strains examined in this study. In that study, MA lines were propagated on LB agar, and mutations accumulated during the MA experiment were identified by sequencing the lines at the endpoint. These mutations were subsequently used to estimate the mutation rate and characterize various mutation biases.

While re-examining the mutation data (Datasheet S6) (5), we identified instances in which two or more independent MA lines from the same strain shared either an identical complete set of mutations or a subset of mutations, raising the possibility that these mutations did not arise independently during the MA experiment. We considered two potential explanations for this pattern. First, some mutations may have been present at low frequency in the ancestral colony from which the MA experiment was initiated. If such pre-existing variants were carried into multiple MA lineages during the establishment of the experiment, the same mutation(s) would subsequently be detected in several independent lines. In this scenario, the shared mutations would represent ancestral standing variation rather than independent mutations that arose during the MA experiment. Second, the overlap could have resulted from cross-contamination between MA lineages during the course of the experiment. A contamination event between two lineages could result in the transfer of a lineage carrying previously accumulated mutations into another MA lineage, producing identical or partially overlapping mutation sets. Such mutations would likewise violate the assumption that mutations observed in different MA lines represent independent mutational events. Because the available data do not allow us to unambiguously distinguish between these two possibilities, we took a conservative approach to minimize the potential influence of non-independent mutations on our estimates.

For analyses of mutation biases, we retained only unique mutation sets for each strain. When the exact same set of mutations was observed in multiple independent MA lineages, the mutation set was counted only once rather than treating each occurrence as an independent observation (Datasheet S7). Thus, duplicated mutation sets were removed from the dataset, preventing potentially shared ancestral or contamination-derived mutations from being disproportionately represented in estimates of mutation spectra and other mutation biases. Table S1 shows corrected mutation biases.

For mutation rate calculations, we applied a more stringent correction. When mutations were shared between MA lineages, the shared mutations were removed from both lineages rather than assigning the mutation to either lineage or counting one occurrence as independent (Datasheet S8). The number of mutations used for calculating the mutation rate was therefore based only on mutations that could be considered unique to a given MA lineage. The corrected number of mutations was then used to recalculate the mutation rate for each strain (Table S1). Following (6) we calculated the mutation rate of each strain using the following formulae. $m$ = mutations per MA lineage, $g$ = generations per MA lineage; mutation rate of an MA line, $r = m / g$. The mean overall mutation rate was calculated as: $R = M / G$, where $M = \Sigma m$ and $G = \Sigma g$ over all MA lines, including lineages in which no mutations were observed. The variance of R was estimated as: $Var = [\ \Sigma(R - r)^2 / (N - 1)\ ] \times [\ \Sigma g^2 / G^2\ ]$, where N is the number of MA lines analysed for that strain. The 95% confidence interval was then calculated as: 95% CI = $R \pm (\sqrt{Var} \times t_{0.05,\ N-1})$, where $t_{0.05,\ N-1}$ is the two tailed critical value of Student's t-distribution at a significance level of 0.05 with N-1 degrees of freedom. The per-nucleotide mutation rate and its confidence interval were obtained by dividing R and the CI half-width by the genome size.

#### Testing the effect of $S_b$ on adaptation rate

From Experiment 1, to test the effect of $S_b$ on adaptation rate, we analyzed the quantitative distribution of adaptation rate differences between high-$S_b$ and low-$S_b$ strains. We calculated adaptation rate differences for all possible pairs of high-$S_b$ and low-$S_b$ strains within each environment, as the difference in the mean percent growth rate increase $[((final\ growth\ rate - initial\ growth\ rate)/\ (initial\ growth\ rate)) \times 100]$ of each strain across all replicate populations during evolution. Because mutation rate varies more substantially than the fraction of beneficial mutations ($f_b$) among our strains, we expected mutation rate to have a stronger influence on $S_b$. This led to the predicted overall rank order of strains: *ΔmutT* > *ΔmutH* > *ΔmutY* > *Δnth-nei* > WT. Based on this ranking, there are 10 possible high-$S_b$ versus low-$S_b$ strain pairs in each environment (60 pairs in total across 6 environments). We tested the effect of environment on adaptation rate differences using a one-way ANOVA. Environment had no significant effect on adaptation rate differences at day 16 (one-way ANOVA: ($F = 0.57$, $p = 0.72$). In contrast, environment significantly affected adaptation rate differences at day 48 ($F = 5.45$, $p = 3.95e-4$). Post hoc Tukey's HSD tests showed that adaptation rate differences in gluconate were significantly lower than other environments. No other pairwise comparisons were statistically significant, hence we compiled the pairwise comparisons across environments, leading to 60 pairs in total across 6 environments. The resulting distribution of adaptation rate differences provides a direct and quantitative measure of the effect of $S_b$ on adaptation rate. We tested whether this distribution was significantly shifted right (relative to zero) using a one-sample, one-tailed t-test. A significant right shift would indicate that high-$S_b$ strains tend to adapt faster than low-$S_b$ strains, consistent with our prediction. Similarly, for Experiment 2, we tested the effect of mutation bias by performing all possible pairwise comparisons between the 12 Δ*mutT* and 12 Δ*mutH* clones, yielding a total of 144 pairwise comparisons. For each clone pair, we calculated the adaptation rate difference as the difference between the mean percent growth rate increase of the Δ*mutT* clone and that of the Δ*mutH* clone, where the mean for each clone was calculated across all five replicate populations.

#### Quantifying variation in adaptation rate across replicate populations

In Experiment 1, we quantified variation in adaptation rate across replicate populations by calculating adaptation rate differences between all unique pairs of independently evolved biological replicates of the same strain in the same environment. Pairwise comparisons were generated without replacement in a fixed order (1–2, 1–3, …, 1–6, 2–3, …, 5–6), such that each replicate pair was included only once. This yielded 15 pairwise comparisons for each strain–environment combination at day 16 (6 replicates) and 3 pairwise comparisons at day 48 (3 replicates). Across all five strains and six environments, this resulted in a total of 450 pairwise comparisons at day 16 and 90 pairwise comparisons at day 48.

In Experiment 2, we quantified variation in adaptation rate among the five independently evolved replicate populations derived from each founder clone. For each of the 12 founder clones of a given strain, all unique pairwise comparisons among the five replicate populations were generated without replacement, yielding 10 replicate pairs per clone and 120 pairwise comparisons per strain.

### Identifying high-frequency mutations

For each independently evolved population, mutations were grouped into frequency bins (0-10%, 10-20%, …, 90-100%) based on their observed frequency. We defined mutations with a frequency of ≥30% as high frequency. Because the mutation frequency spectrum was strongly skewed toward rare mutations (Fig. S6), thresholds higher than 30% resulted in many strain–environment combinations with no high-frequency mutations. We therefore selected 30% as the highest threshold that yielded a non-zero average number of high-frequency mutations per strain-environment combination, enabling meaningful comparisons of the effects of mutation rate and mutation bias on the accumulation of high-frequency mutations.

### Calculating mutation and gene level repeatability

For each strain–environment combination, we determined the extent of mutational parallelism by classifying each exact mutation or each mutated gene according to the number of independently evolved replicate populations in which it was observed. Mutations or genes were categorized as unique (present in a single replicate population) or shared among 2, 3, 4, 5, or all 6 replicate populations. To quantify the overall level of parallel evolution, we then calculated the total number of mutations (or genes with mutations) that were shared across two or more replicate populations.

### Calculating Jaccard similarity index

For every pair of independently evolved populations within a given strain–environment combination, we quantified genomic similarity using the Jaccard index. For gene-level analyses, the Jaccard index was calculated as the number of genes mutated in both populations divided by the total number of genes mutated in either population. The same analysis was performed at the mutation level using the sets of exact mutations present in each population. All unique pairwise comparisons were considered, yielding 15 pairwise comparisons for each strain–environment combination at day 16 (6 replicate populations) and 3 pairwise comparisons at day 48 (3 replicate populations).

### Estimating the distribution of selection coefficients

To estimate the distribution of selection coefficients (*s*), we analysed changes in allele frequencies (*p*) over time (*t*) using the logistic model, $\frac{dp}{dt} = sp(1-p)$. Because sequencing data were available at both day 16 and day 48 only for replicate populations 1, 2, and 6, the analysis was restricted to these replicate populations. For each evolved population, allele frequencies at the two sampling time points were converted from percentages to proportions by dividing by 100. To avoid numerical instability during the logit transformation, allele frequencies exactly 0 or 1 were converted to 0.00001 and 0.99999 respectively. The initial ($p_i$) and final ($p_f$) allele frequencies were then used to estimate the selection coefficient. Assuming a time interval of approximately 200 generations (32 days) between the two timepoints, the selection coefficient was calculated as

$$s = \frac{\ln\left(\frac{p_f}{1-p_f}\right) - \ln\left(\frac{p_i}{1-p_i}\right)}{\Delta t}$$

### Packages used

Data processing and statistical analyses were performed in Python version 3.11.4 (7) using pandas (8), NumPy (9), SciPy (10), and the standard os and itertools libraries. Additional statistical analyses and data visualization were performed in R version 4.5.0 (11) using packages ggplot2 (12), ggpubr (13), dplyr (14), tidyr (15), lme4 (16), lmerTest (17), MAAS (18) and pheatmap (19). Final figure assembly and graphical editing were carried out using Adobe Illustrator (Adobe Inc.) (20).

**SUPPLEMENTARY FIGURES**

**Figure S1: Adaptation over time. (A)** Spearman's rank correlation between % growth rate change from days 0–16 and days 16–48. Growth rate change was calculated as: $[(Final\ growth\ rate - Initial\ growth\ rate)\ /\ Initial\ growth\ rate)\ \times\ 100]$. **(B)** Comparison of the percent change in growth rate per day from day 0 to 16 and from day 16 to 48 (paired t-test). Lines connect means for each strain in each environment. **(C)** Same as panel (B), but excluding galactose populations to allow better visualization of results in other environments. In panels A–C, each point represents the mean ± SE across biological replicates of a strain evolving in an environment (n=6 at day 0 and day 16; n=3 at day 48). **(D)** Change in population size (the number of colony forming units, CFU/mL) of each strain over 48 days of laboratory evolution in galactose and gluconate (paired t-test). These environments were chosen for this assay because they showed the highest and lowest adaptation rate respectively. Each point represents mean ± SE across technical replicates (n=2-3).

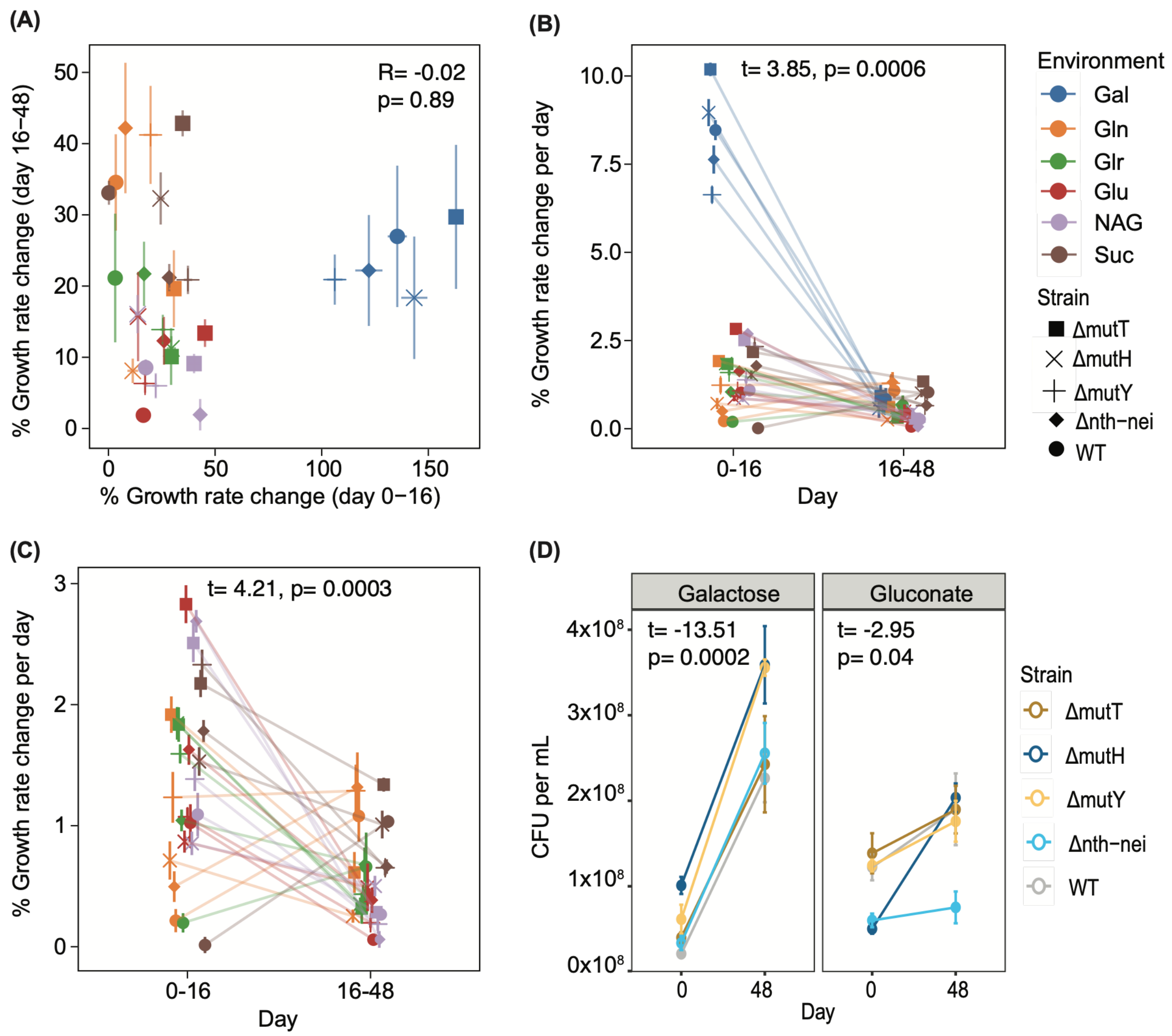

**Figure S2:** Spearman's rank correlation between $S_b$ (estimated as the product of mutation rate and $f_b$ in glucose (3), shown on a log scale for better visualization) and adaptation rate in glucose. Data include six replicate populations per strain at day 16 and three replicate populations per strain at day 48.

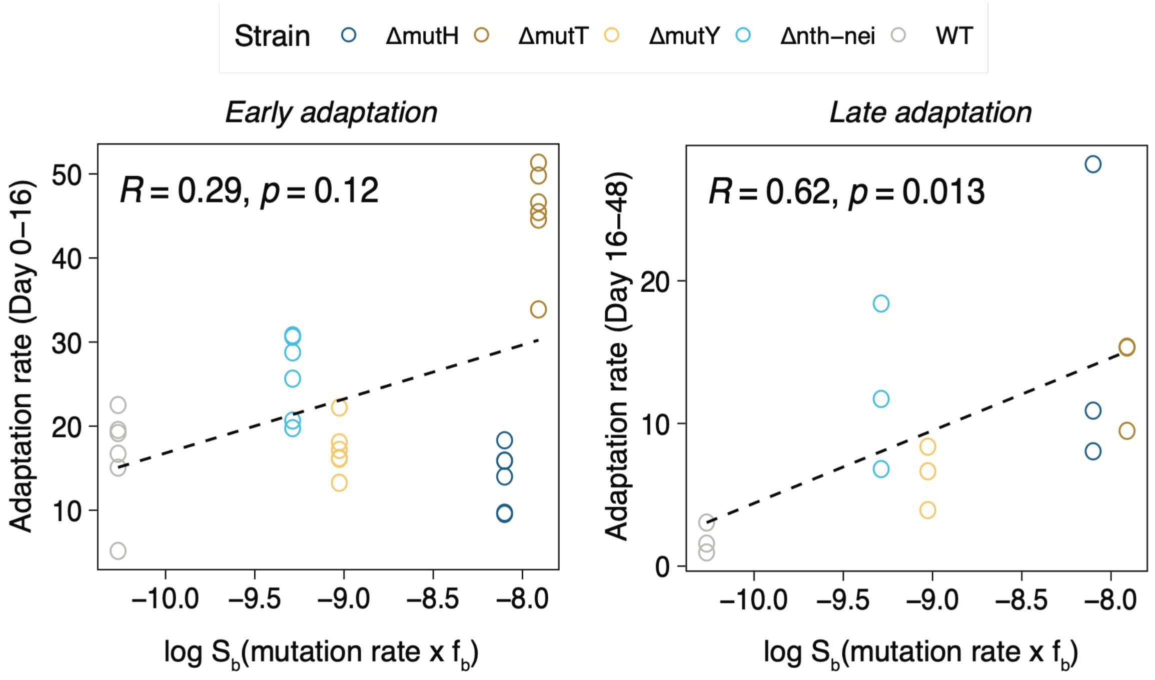

**Figure S3:** For each strain, adaptation rate differences across biological replicates (Fig. 2C) were pooled across environments, and the variance of this pooled distribution was calculated. An F-test (function “var.test” in R) was used to determine whether the variance differed significantly between strains with reversed bias and reinforced bias, for a given mutation rate category.

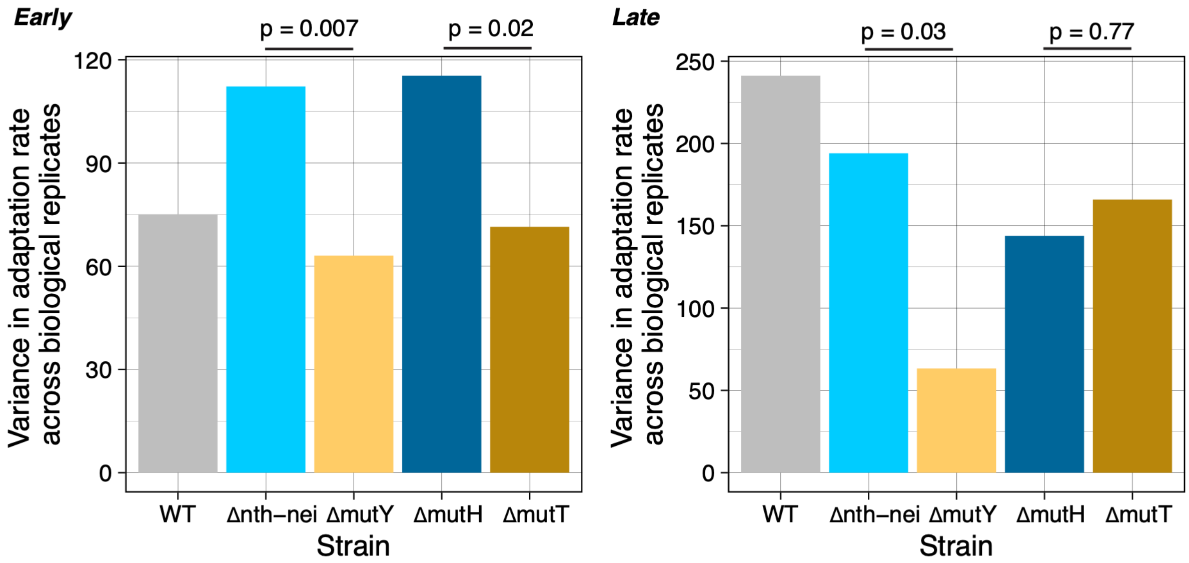

**Figure S4. Effect of population size on the rate of adaptation.** Spearman's rank correlation between the ratio of population size of the High-$S_b$ strain (strain A) and the Low-$S_b$ strain (strain B), as estimated from colony-forming units (CFU/mL), and the difference in their rates of adaptation. Each point represents a strain pair in a single environment (60 pairs in total across six environments).

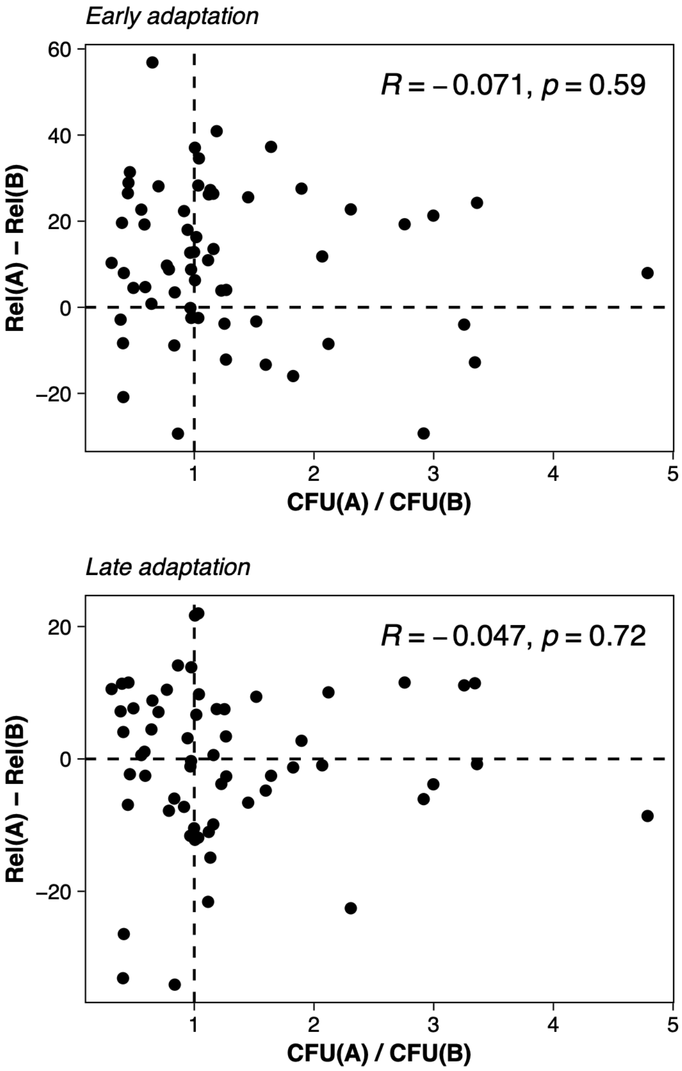

**Figure S5. Effect of initial fitness on the rate of adaptation.** Spearman's rank correlation between the initial fitness and adaptation rate of a population. Data include six replicate populations per strain at day 16 and three replicate populations per strain at day 48. Dashed lines show the best-fit linear regression.

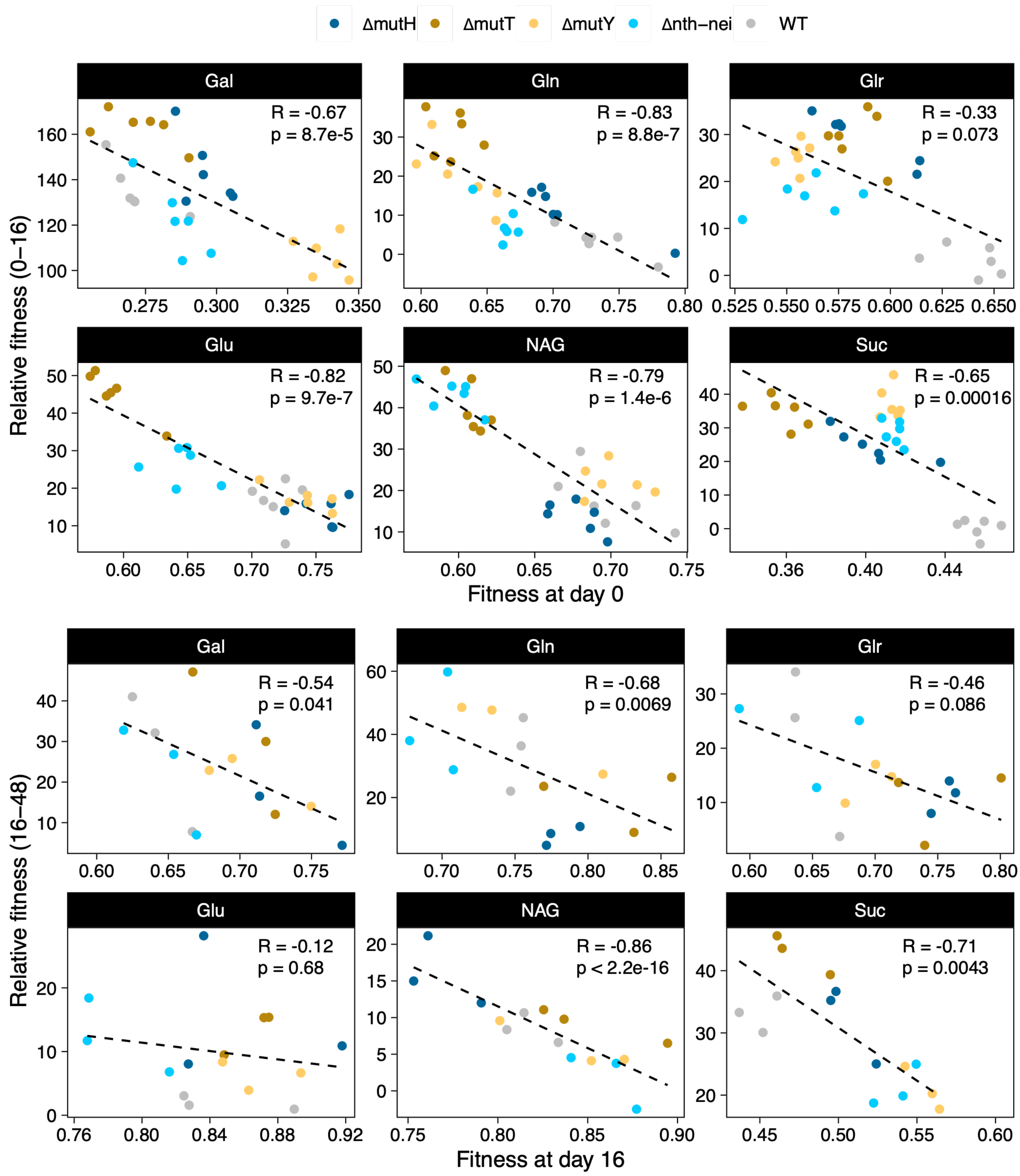

**Figure S6: Initial growth rate of each strain in different environments** (mean ± SD, n=6 biological replicates). Different letters indicate statistically significant differences between strains (paired t-tests, $p < 0.05$).

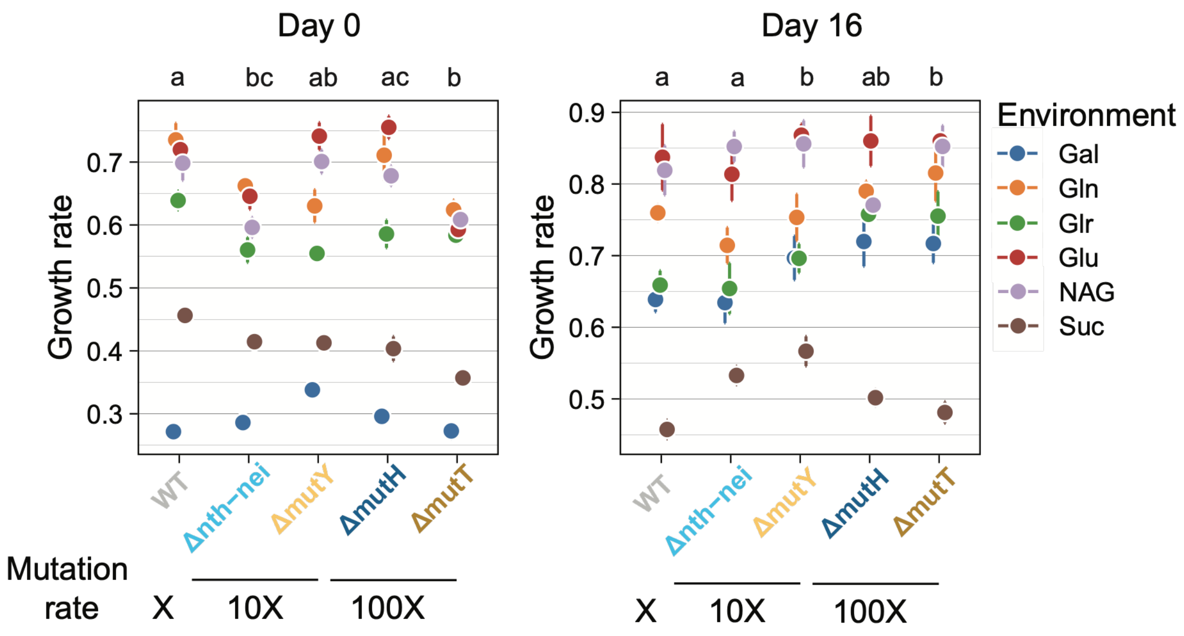

**Figure S7: Effect of initial fitness on adaptation rate**. Spearman's rank correlation between initial fitness and adaptation rate across evolved populations. The analysis include six independently evolved replicate populations per strain per environment at day 16 and three independently evolved replicate populations per strain per environment at day 48.

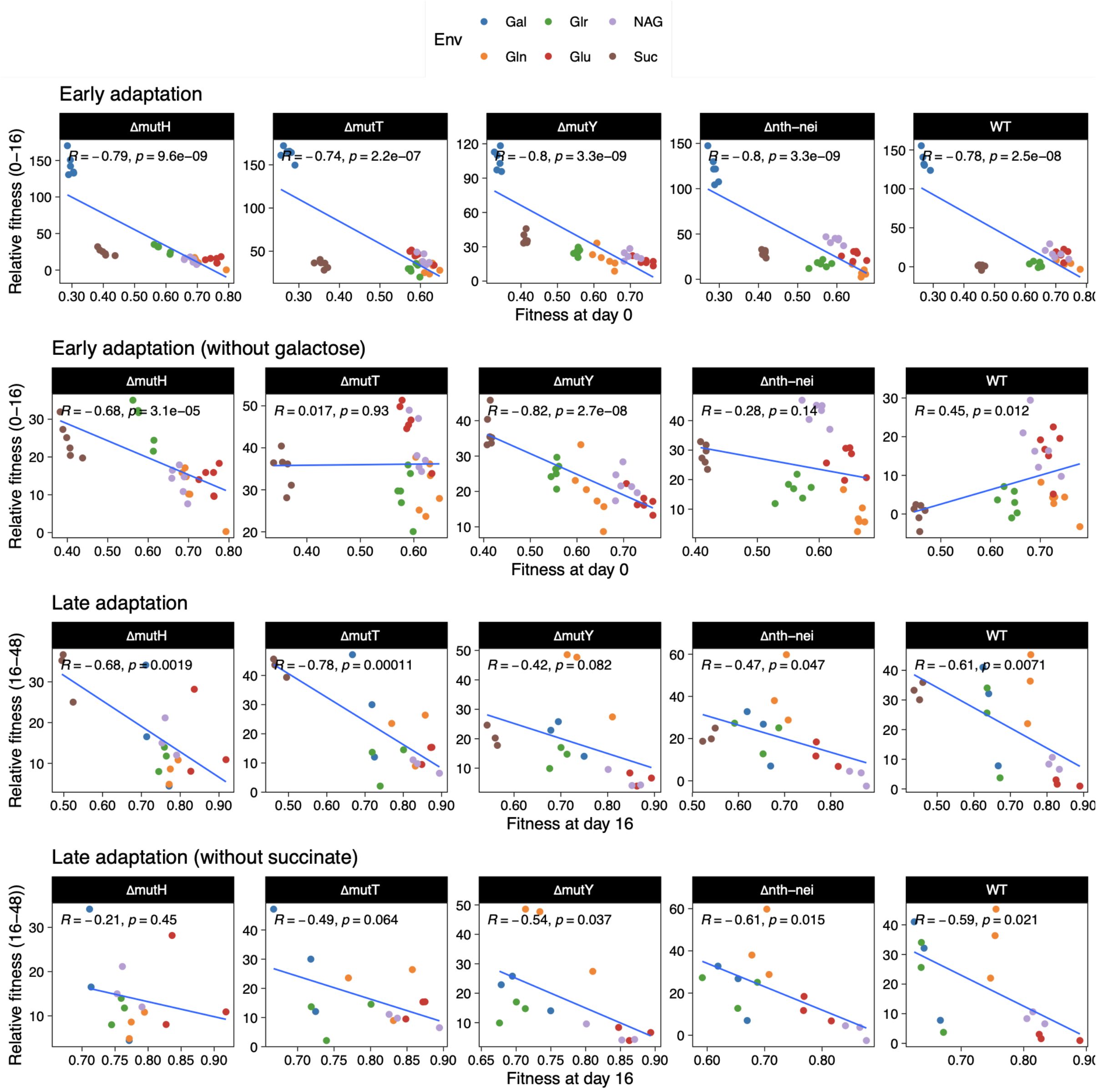

**Figure S8: Effect of initial fitness on adaptation rate in Experiment 2.** Spearman's rank correlation between initial fitness and relative fitness of evolved *ΔmutT* and *ΔmutH* clones. Each point represents the mean ± SD of the initial fitness (across 3–6 technical replicates) and the relative fitness (across five independently evolved biological replicate populations). Lines indicate linear regression fits for significant correlations (only significant for *ΔmutT*); solid line: all data, dashed line: excluding the outlier identified using the 1.5x interquartile range criterion. Coefficients from a linear regression (model: Relative fitness at day 16 ~ Fitness at day 0 x Strain); estimates ± SE: intercept = 55.42 ± 37.56 (t = 1.48, p = 0.16); Fitness at day 0 = −56.87 ± 52.27 (t = −1.09, p = 0.29); Strain = 22.35 ± 44.73 (t = 0.50, p = 0.62); Fitness at day 0 x Strain = −25.52 ± 62.57 (t = −0.41, p = 0.69). Model fit: F-statistic = 3.19, p = 0.046, $R^2$ = 0.324, adjusted $R^2$ = 0.222.

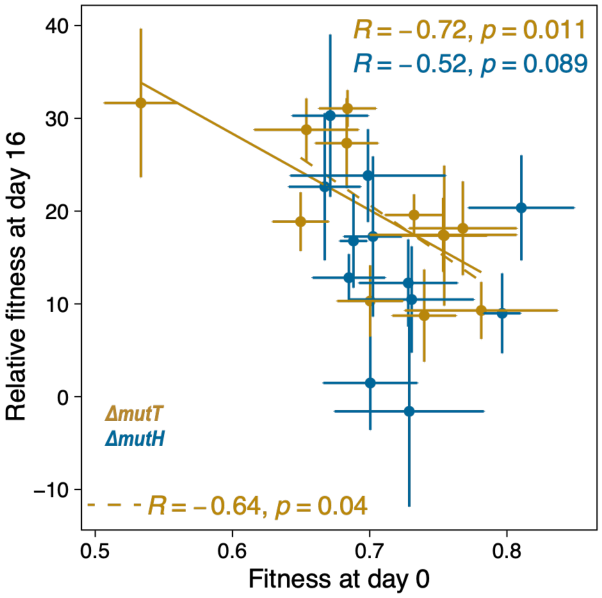

**Figure S9. Total mutations observed as a function of mutation rate.** Spearman rank correlation between the total number of mutations per population and baseline mutation rate of the strain (corrected estimates from mutation accumulation lines propagated on LB agar from (3), as described in the supplementary methods. Mutation rates did not differ significantly between Δ*mutT* and Δ*mutH*, or between Δ*mutY* and Δ*nth-nei* (overlapping 95% CIs), so each pair was assigned a tied rank in the Spearman correlation analysis. Each point represents an independently evolved population of a strain in a specific environment, sampled at day 16 (early) or day 48 (late). Also see Table S2 for the output of negative binomial GLMs fit (using R package MASS::glm.nb) to the model accounting for environment effects: total number of mutations ~ mutation rate * environment.

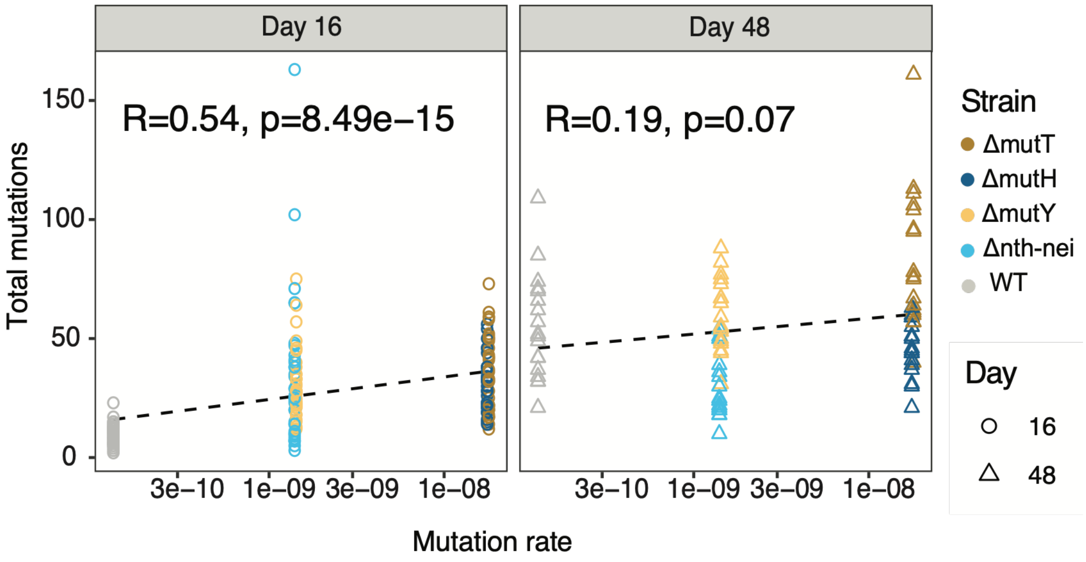

**Figure S10. Frequency distribution (site frequency spectrum) of mutations in evolved populations.** Boxplots show the proportion of mutations observed within each frequency bin. Each point represents an independently evolved population of a strain in a specific environment (n=6 and n=3 per strain per environment at day 16 and 48 respectively). Box boundaries represent the first and third quartiles, the median is indicated as a horizontal line, and whiskers extend to 1.5x the interquartile range. Points beyond whiskers are identified as outliers.

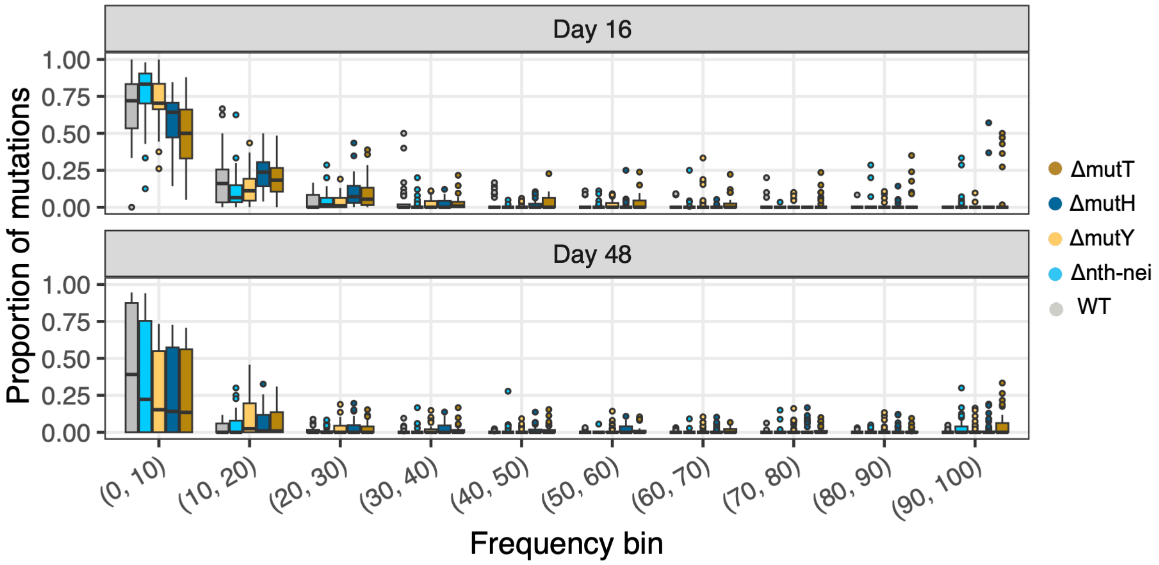

**Figure S11: Genetic repeatability during adaptation, at the level of specific mutations (compare with Fig. 4 for gene level repeatability). (A)** Number of mutations that were observed in at least two independent replicate populations of the same strain evolving in a given environment. **(B)** Pairwise Jaccard similarity index calculated for each strain within each environment. Each point represents the mean ± SD for a given strain across replicate pairs within each environment. At day 16, six replicate populations per strain yield 15 pairwise comparisons, whereas at day 48, three replicates yield three pairwise comparisons. Reported p-values were derived from paired t-tests. Jaccard index at the mutation level decreased significantly from day 16 to 48 (paired t-test: t = 3.173, df = 29, p = 0.003; note the distinct y-axes between day 16 and day 48 plots).

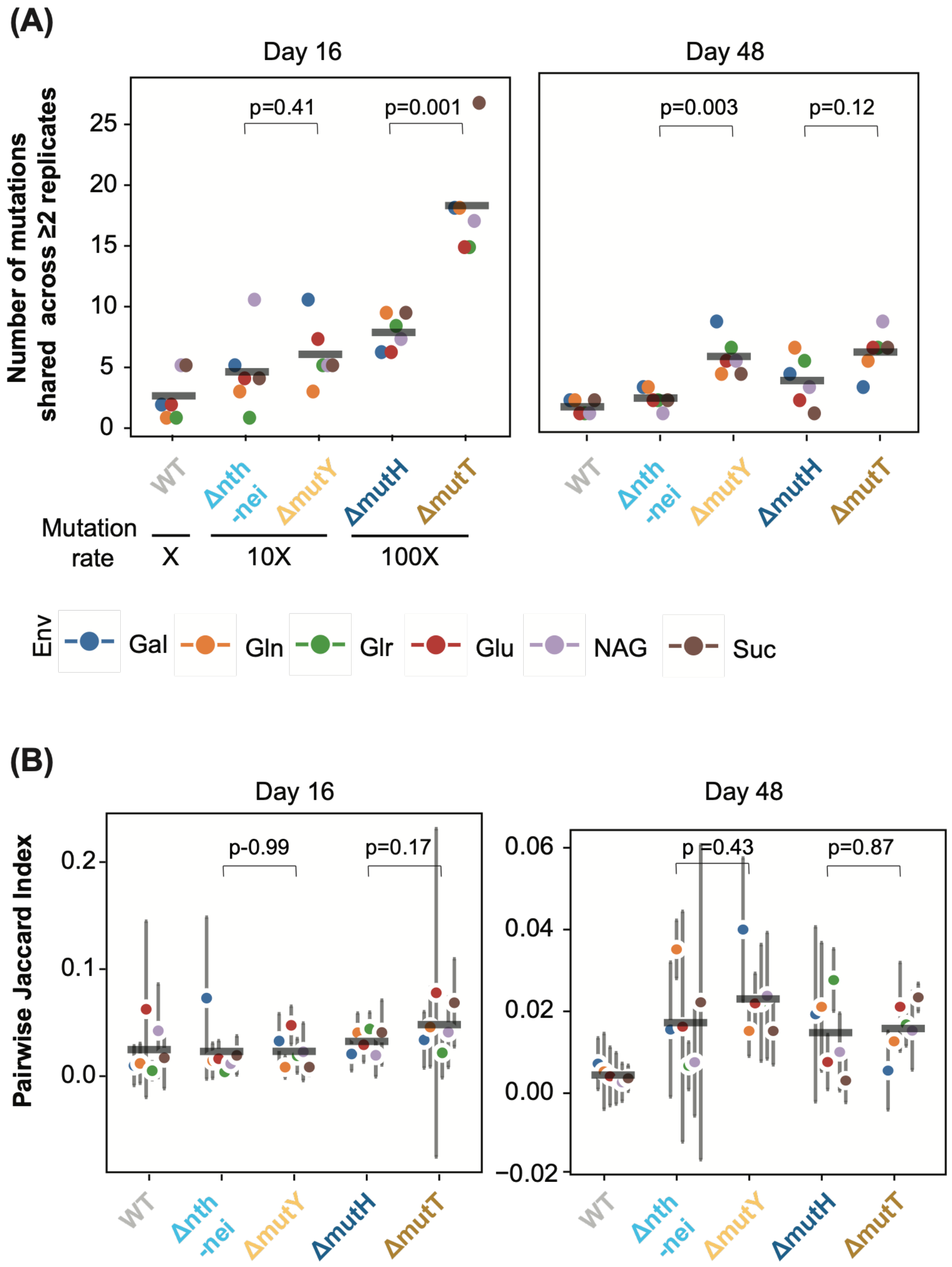

**Figure S12: Estimated selection coefficients for observed mutations in evolved populations. (A)** Distribution of selection coefficients (*s*) of all mutations detected at day 16 in each population, estimated as: $log\left(\frac{freq_{day48}}{1-freq_{day48}}\right) - log\left(\frac{freq_{day16}}{1-freq_{day16}}\right) / (Number\ of\ generations)$. The analysis was restricted to replicates that were propagated to day 48 (replicates 1, 2, and 6); data were pooled across replicates and environments for each strain. Mutation frequencies of 1 and 0 were adjusted to 0.999999 and 0.000001, respectively, to avoid infinite values. The solid black line indicates the median *s* for each strain. Numbers in parentheses in the top right and top left corners indicate the fractions of beneficial (*s* > 0) and deleterious (*s* < 0) mutations respectively. **(B)** Distribution of selection coefficients for beneficial mutations only; the dotted and solid line marks the mean and median respectively (for beneficial mutations only) for each strain.

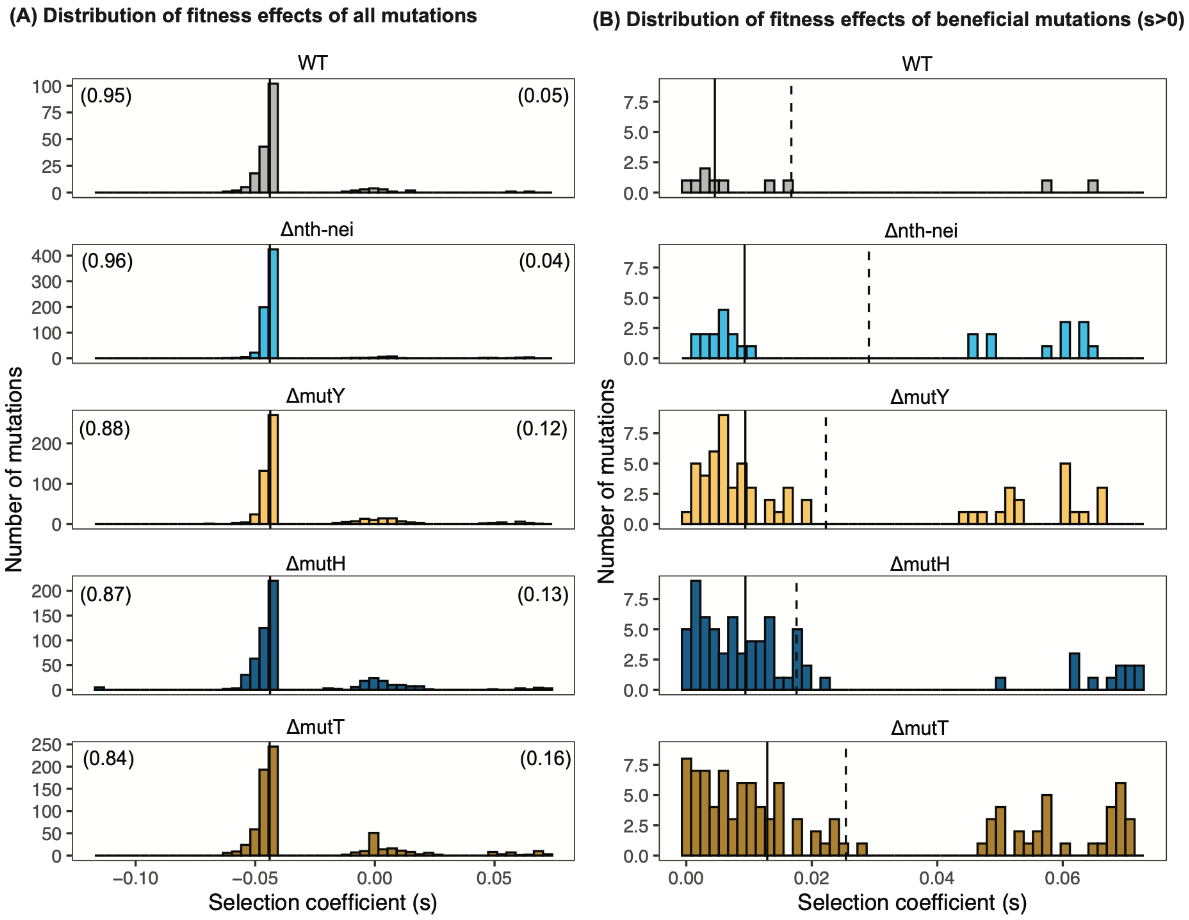

**Figure S13: Correlation between mutation bias under mutation accumulation (MA) and selection.** **(A)** Correlation between the transversion bias (Tv bias) observed under genetic drift (Table S1, see supplementary materials and methods for corrected estimates of mutation bias under drift following (3)) and under selection (evolved populations from the current study). Each point represents the mean ± SD across all environments for a given strain. Circles indicate measurements from day 16 (early phase), and triangles indicate those from day 48 (late phase) of adaptation. Tv bias is calculated as Tv / (Tv + Ts), where Tv and Ts represent the number of transversions and transitions, respectively. **(B-E)** Other mutational biases in evolved populations, compared to MA. **(B)** SNP bias, calculated as SNP / (SNP + Indel). **(C)** GC→AT bias, calculated as the number of GC→AT / (GC→AT + AT→GC) mutations. **(D)** Non-synonymous bias, calculated as the number of Non-synonymous / (Non-synonymous + Synonymous) mutations. **(E)** Coding bias, calculated as the number of Coding / (Coding + Non-coding) mutations. In panels A-E, Spearman's rank correlation is shown, and diagonals represent a slope of 1, i.e., when the bias under MA and selection is identical. **(F)** Single nucleotide polymorphism (SNP) spectrum under MA (leftmost bar in each panel) vs. under selection after 16 and 48 days of evolution.

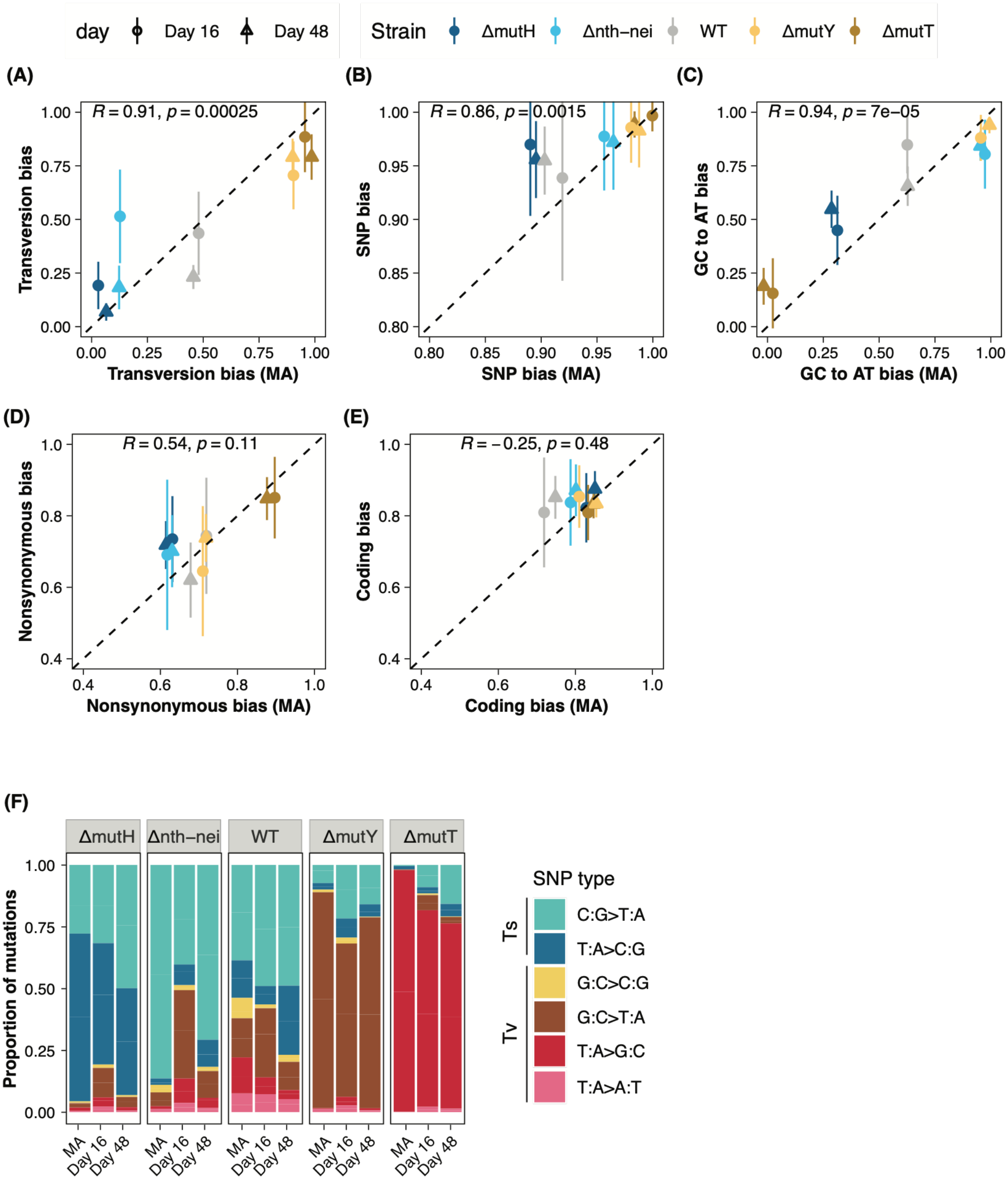
day
Day 16
Day 48
Strain
ΔmutH
Δnth−nei
WT
ΔmutY
ΔmutT
(A)
R = 0.91, p = 0.00025
Transversion bias
Transversion bias (MA)
(B)
R = 0.86, p = 0.0015
SNP bias
SNP bias (MA)
(C)
R = 0.94, p = 7e−05
GC to AT bias
GC to AT bias (MA)
(D)
R = 0.54, p = 0.11
Nonsynonymous bias
Nonsynonymous bias (MA)
(E)
R = −0.25, p = 0.48
Coding bias
Coding bias (MA)
(F)
Proportion of mutations
MA
Day 16
Day 48
SNP type
Ts
Tv
C:G>T:A
T:A>C:G
G:C>C:G
G:C>T:A
T:A>G:C
T:A>A:T

**Figure S14: Mutation biases at the most frequently mutated loci.** The heatmap shows the most frequently mutated loci across strains, environments, and time points. Only genes mutated in at least two replicate populations of a strain and independently mutated across at least two strains, environments, or timepoints are included. The grayscale intensity indicates the proportion of replicates in which a given gene was mutated, for each strain and environment at each time point. The top three most commonly mutated loci are marked in red, and are also shown in Figure 5B.

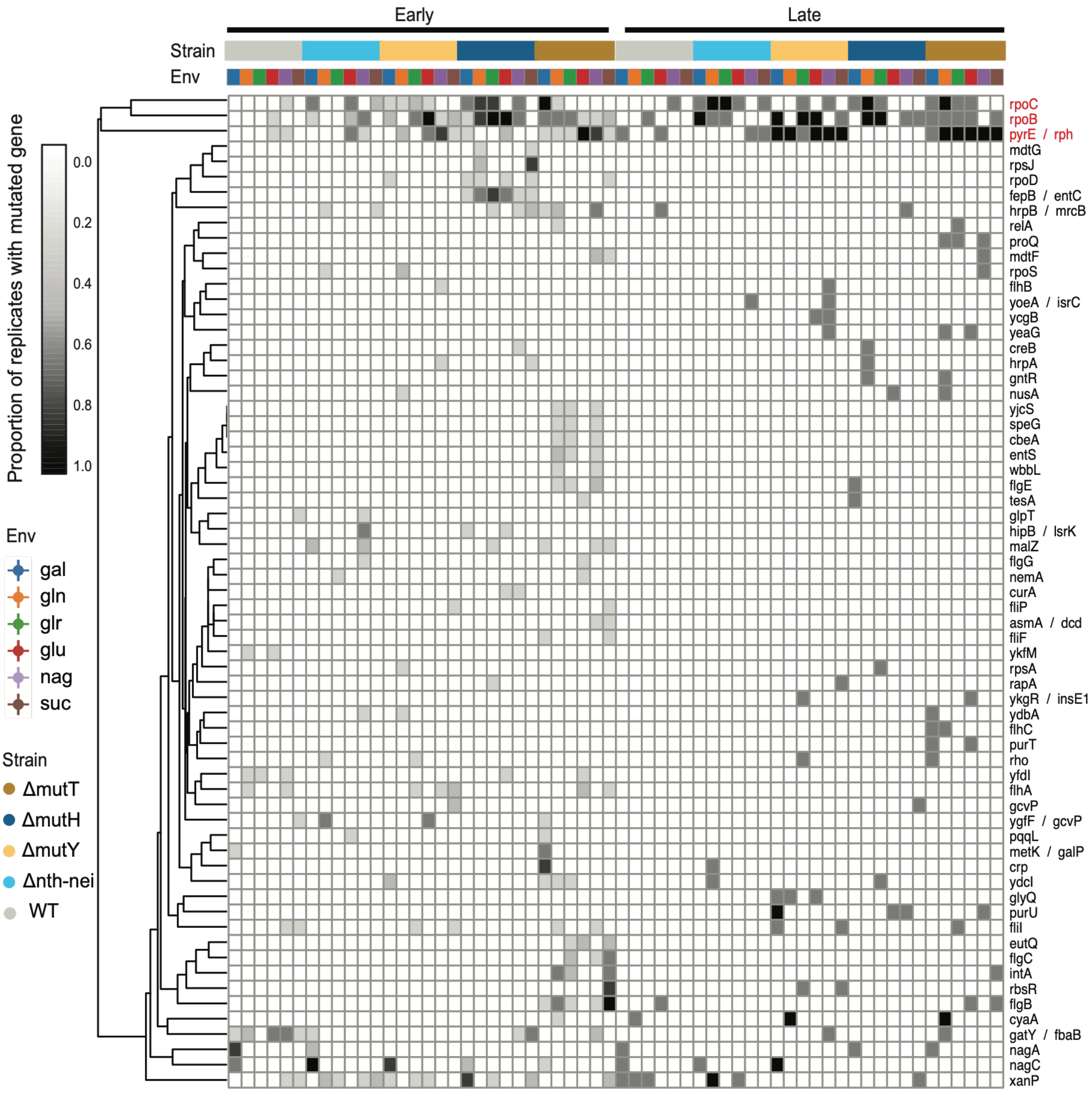

**Figure S15: Fitness effects of observed amino acid changes in *rpoB*.** The relative fitness of 11 amino acid substitutions in *rpoB* found in our dataset that were reported in a prior study (21). Relative fitness was measured using bulk competitions and deep mutational scanning (DMS), tested under five conditions: M9 minimal media supplemented with glucose (Glu), glycerol (Gly), galactose (Gal), glucose with high osmolarity (Nac), and glucose with butanol (But). The plot shows the distribution of fitness effects for substitutions observed in each of our strains, pooled across replicates and environments in our study (n indicates the total number of amino acid changes x number of environments its fitness was measured i.e., 5 for each strain).

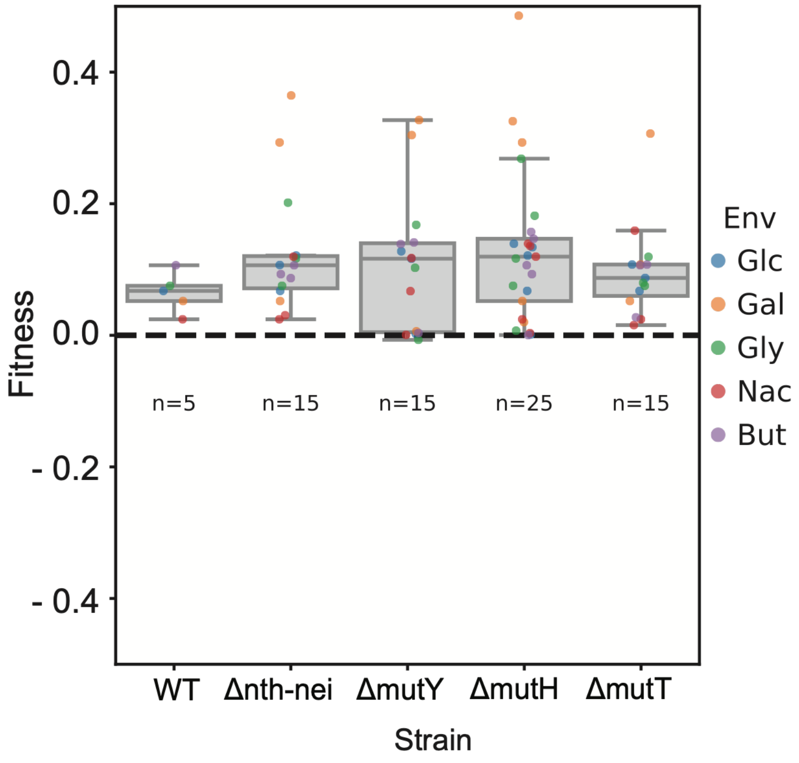

**Figure S16: The location of mutations observed in the *pyrE/rph* intergenic region**. The schematic shows the focal genomic region (numbers in parentheses indicate the start and end of the respective gene). The x-axis represents the position upstream of the *pyrE* gene. The ancestral deletion causing a frameshift (found in all our strains) at the C-terminal end of *rph* is indicated. Each circle and triangle indicates the position of a SNP or indel observed in our evolved populations.

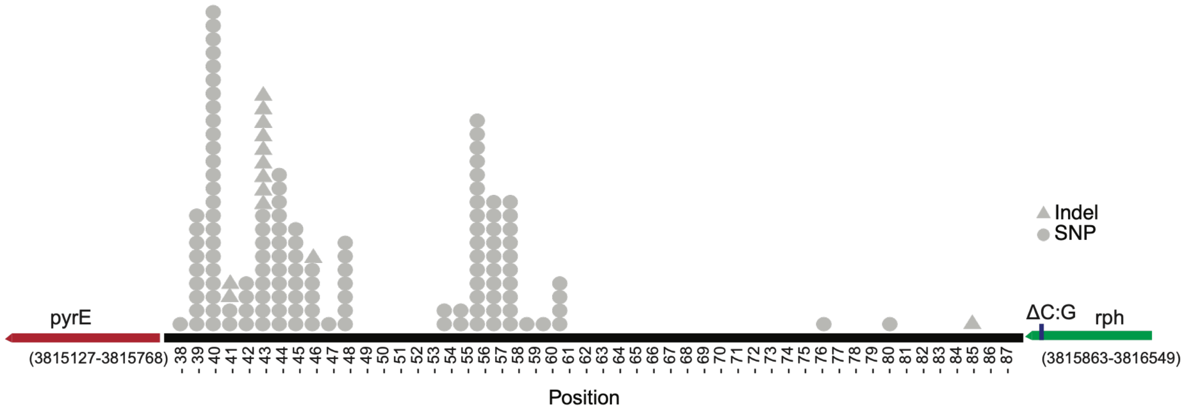

**Figure S17. Frequency and distribution of mutations at the most frequently mutated loci. (A)** Allele frequency of mutations in *rpoB*, *rpoC*, and *pyrE/rph* loci. Each point represents the frequency of a single mutation observed in an evolved population, at a specific time. Data are pooled across environments and replicate populations for each strain. **(B)** Density plots showing the distribution of mutation positions within the three loci of interest, compiled across days and environments for each strain. For *rpoB* and *rpoC*, 0 at the x-axis indicates the start position of the gene. For *pyrE/rph*, the two solid black lines represent the end of *pyrE* and the start of the *rph* gene respectively (see Figure S9 for a schematic of the locus). For *rpoB* and *rpoC*, mutation distributions differed significantly between Δ*mutH* and each of the other three mutator strains. For *pyrE/rph*, the distribution in Δ*mutT* differed significantly from those of the other four strains, while WT also differed significantly from Δ*mutH* and Δ*mutY* (Kruskal–Wallis test, $p < 0.05$ for all comparisons).

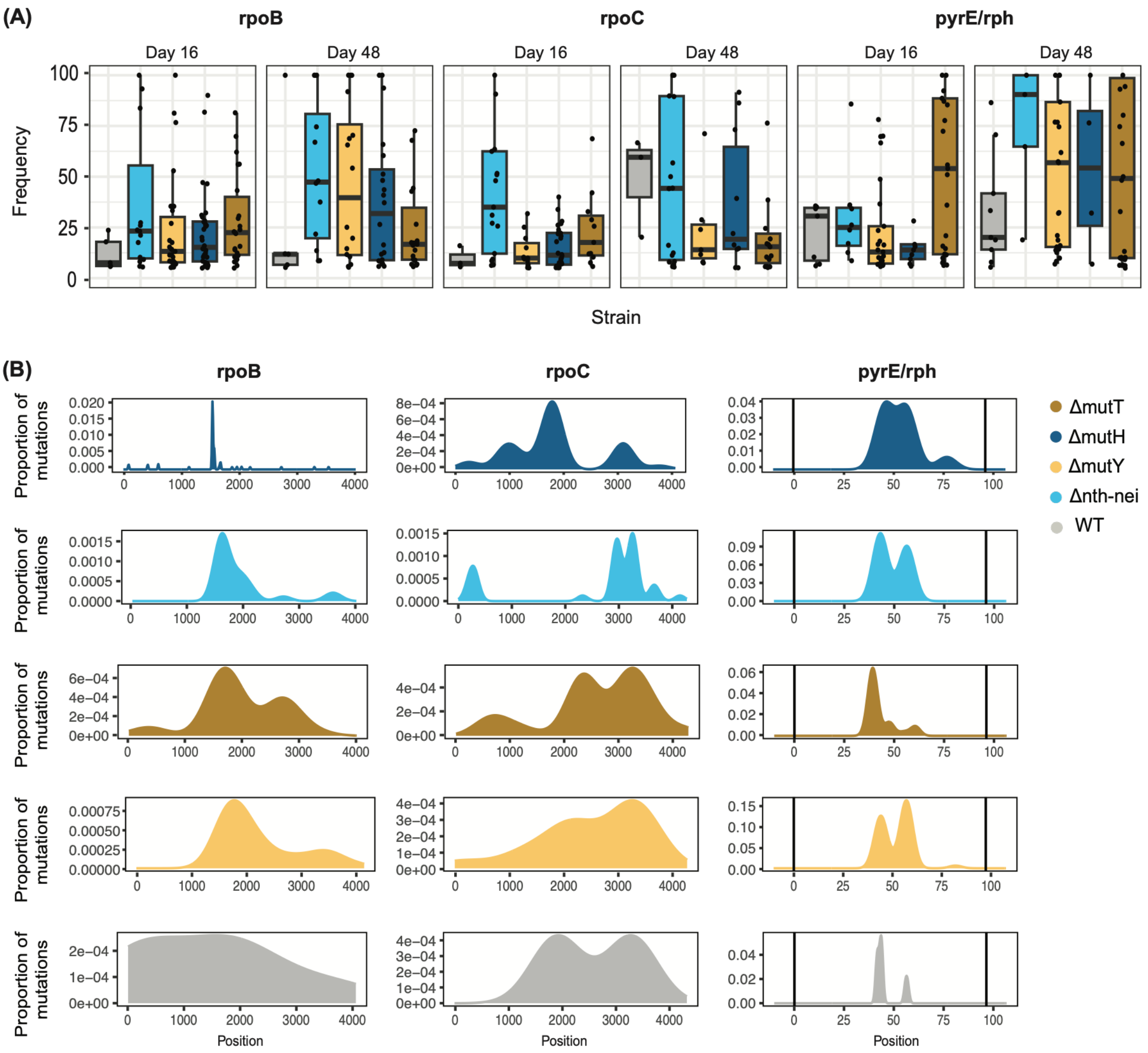

**Figure S18. Types of amino acid changes in *rpoB* and *rpoC* observed across strains.** The heatmap indicates the number of mutations of each type observed for a given strain. Data across replicates and environments have been pooled for each strain. We observed a significant effect of strain identity on the changes in each gene (Chi-square test, $p < 0.05$ in each case).

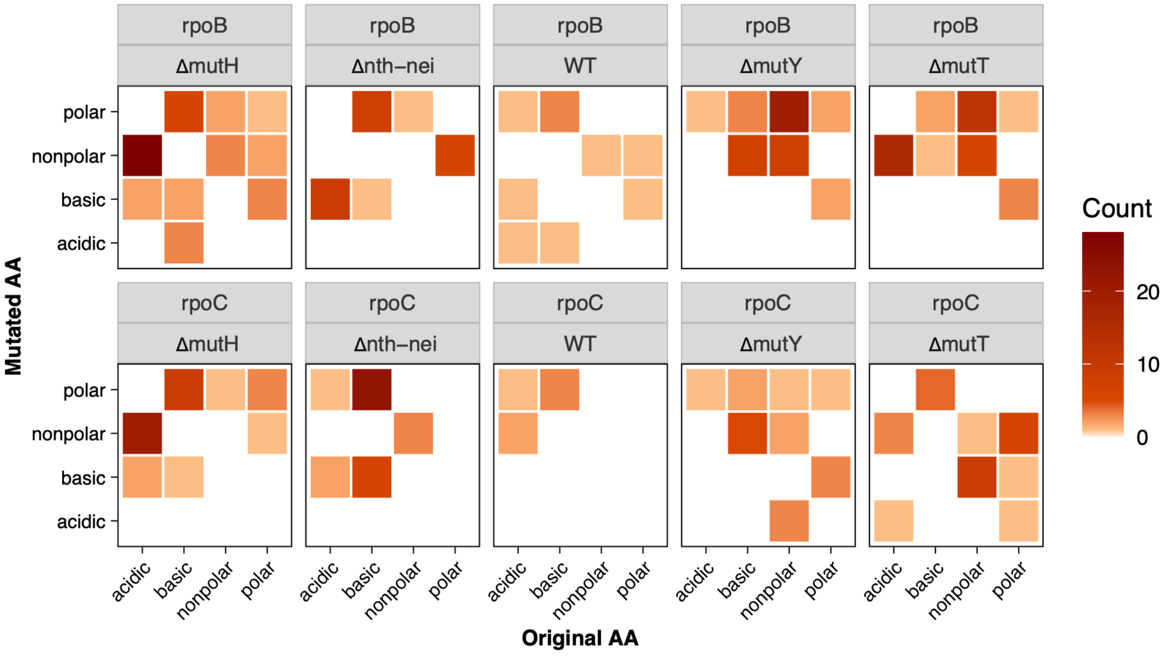

**Figure S19:** Mutation spectra across strains for genes mutated in multiple replicates of a given strain and shared across strains, while remaining specific to a particular environment. The color key on the left indicates the most common SNP types observed in MA, serving as a reference for comparison.

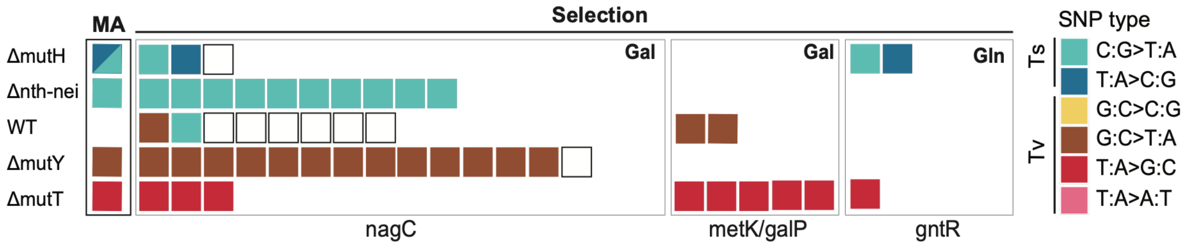

**Figure S20: Growth rate measurements show high repeatability across machines and days. (A)** Spearman's rank correlation for growth rates measured on two different Biotek spectrophotometers. **(B)** Spearman's rank correlation between ancestral growth rate measured along with day 16 evolved populations and day 48 evolved populations across several microplates, on two different Biotek spectrophotometers. **(C)** Difference in the growth rates of the WT reference strain measured in a specific microplate, from the mean value across all microplates (the gray bar represents a difference of ± 0.05).

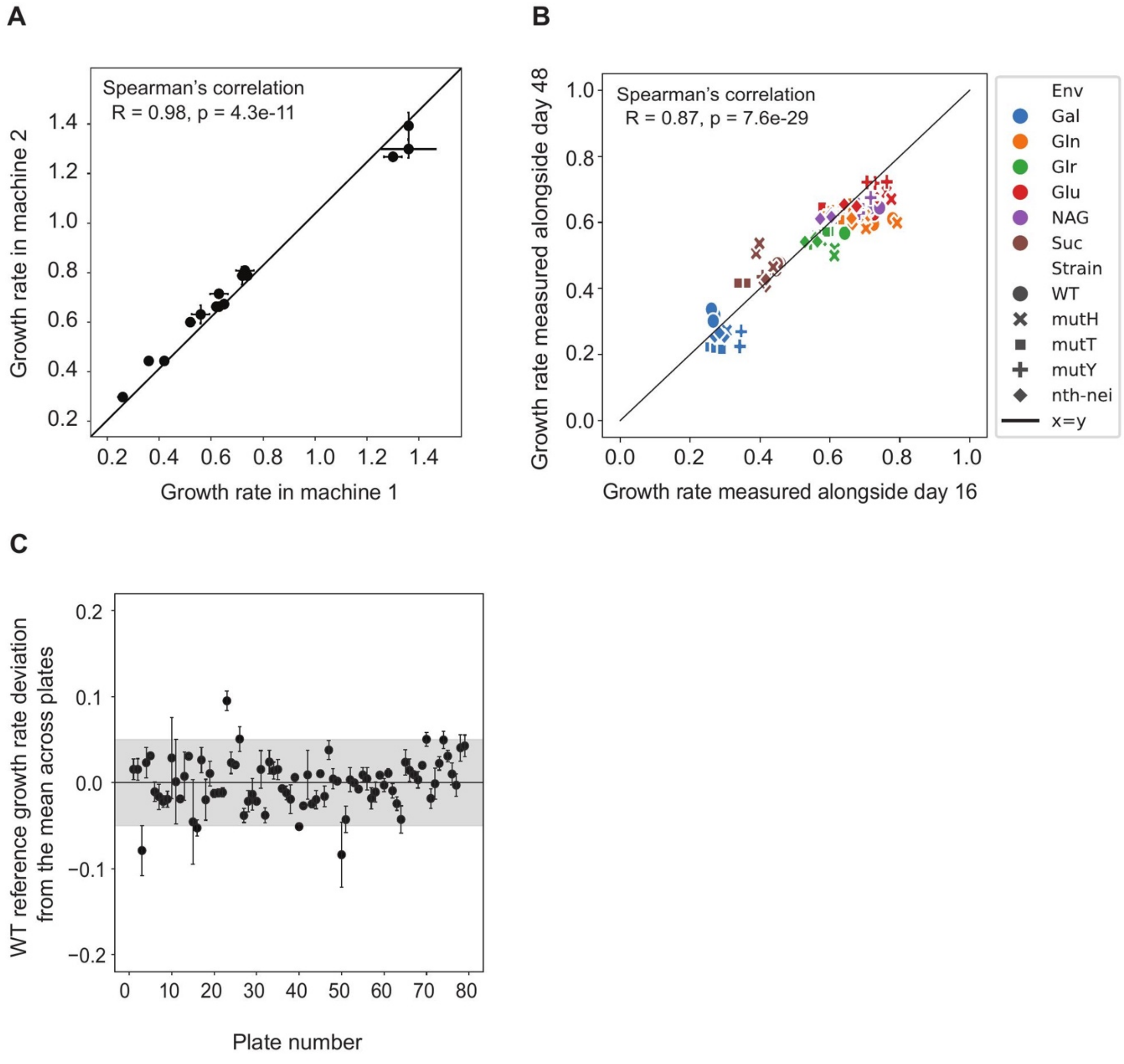

### SUPPLEMENTARY TABLES

**Table S1:** Mutation rates and spectra measured under mutation accumulation (see Supplementary Material and Methods for calculations), and the distribution of fitness effects in M9 minimal media supplemented with 5mM Glucose for the strains used in this study. Data from Sane et al., 2025 (3).

| Strain | WT | ∆nth-nei | ∆mutY | ∆mutH | ∆mutT |
|---|---|---|---|---|---|
| Mutation rate | 1.29e-10 | 1.62e-09 | 1.43e-09 | 1.76e-08 | 1.80e-08 |
| ±95% CI | 3.56e-11 | 1.49e-10 | 1.68e-10 | 1.36e-09 | 1.69e-09 |
| Indel bias | 0.09 | 0.04 | 0.01 | 0.11 | 0.01 |
| ±95% CI | 0.03 | 0.02 | 0.01 | 0.02 | 0.01 |
| Coding bias | 0.72 | 0.80 | 0.84 | 0.84 | 0.85 |
| ±95% CI | 0.05 | 0.04 | 0.03 | 0.02 | 0.03 |
| Nonsynonymous bias | 0.70 | 0.63 | 0.72 | 0.64 | 0.89 |
| ±95% CI | 0.05 | 0.04 | 0.03 | 0.03 | 0.02 |
| Transversion bias | 0.46 | 0.11 | 0.90 | 0.04 | 0.98 |
| ±95% CI | 0.05 | 0.03 | 0.02 | 0.01 | 0.01 |
| GC-->AT bias | 0.65 | 0.96 | 0.97 | 0.30 | 0.01 |
| ±95% CI | 0.05 | 0.02 | 0.01 | 0.03 | 0.01 |
| Fraction of beneficial mutations in M9+Glucose | 0.42 | 0.37 | 0.66 | 0.45 | 0.68 |
| Fraction of deleterious mutations in M9+Glucose | 0.30 | 0.18 | 0.10 | 0.41 | 0.07 |
| Fraction of neutral mutations in M9+Glucose | 0.28 | 0.45 | 0.24 | 0.14 | 0.25 |

**Table S2:** Coefficients from negative binomial generalized linear models (MASS::glm.nb; Total mutations ~ Mutation rate × Environment) fit separately for day 16 and day 48. Overdispersion relative to a Poisson model was confirmed prior to fitting (dispersion test, day 16: p = 3.15e-4; day 48: p = 1.11e-08). Significant effects are highlighted in bold.

| Day | term | estimate | std.error | statistic | p.value |
|---|---|---|---|---|---|
| 16 | (Intercept) | 2.85e+00 | 1.57e-01 | 18.16341 | 1.01e-73 |
| 16 | mutation_rate | 5.09e+07 | 1.35e+07 | 3.754311 | **1.74e-04** |
| 16 | Environmentgln | 3.64e-01 | 2.20e-01 | 1.655654 | 9.78e-02 |
| 16 | Environmentglr | 2.31e-01 | 2.20e-01 | 1.046784 | 2.95e-01 |
| 16 | Environmentglu | 1.29e-01 | 2.21e-01 | 0.585545 | 5.58e-01 |
| 16 | Environmentnag | 6.05e-01 | 2.18e-01 | 2.769605 | **5.61e-03** |
| 16 | Environmentsuc | 4.22e-01 | 2.19e-01 | 1.923813 | 5.44e-02 |
| 16 | mutation_rate:Environmentgln | -4.29e+07 | 1.92e+07 | -2.23975 | **2.51e-02** |
| 16 | mutation_rate:Environmentglr | -1.78e+07 | 1.91e+07 | -0.93043 | 3.52e-01 |
| 16 | mutation_rate:Environmentglu | -2.42e+07 | 1.92e+07 | -1.25886 | 2.08e-01 |
| 16 | mutation_rate:Environmentnag | -5.53e+07 | 1.91e+07 | -2.89479 | **3.79e-03** |
| 16 | mutation_rate:Environmentsuc | -3.70e+07 | 1.91e+07 | -1.93357 | 5.32e-02 |
| 48 | (Intercept) | 3.92e+00 | 1.48e-01 | 26.48406 | 1.48e-154 |
| 48 | mutation_rate | 3.39e+06 | 1.31e+07 | 0.259334 | 7.95e-01 |
| 48 | Environmentgln | -8.30e-02 | 2.10e-01 | -0.39605 | 6.92e-01 |
| 48 | Environmentglr | 6.42e-02 | 2.09e-01 | 0.307348 | 7.59e-01 |
| 48 | Environmentglu | -4.15e-01 | 2.12e-01 | -1.95708 | 5.03e-02 |
| 48 | Environmentnag | -2.18e-01 | 2.10e-01 | -1.03481 | 3.01e-01 |
| 48 | Environmentsuc | 1.76e-01 | 2.08e-01 | 0.844743 | 3.98e-01 |
| 48 | mutation_rate:Environmentgln | 2.06e+07 | 1.84e+07 | 1.119866 | 2.63e-01 |
| 48 | mutation_rate:Environmentglr | 7.28e+06 | 1.84e+07 | 0.395181 | 6.93e-01 |
| 48 | mutation_rate:Environmentglu | 2.67e+07 | 1.86e+07 | 1.436072 | 1.51e-01 |
| 48 | mutation_rate:Environmentnag | 3.05e+07 | 1.84e+07 | 1.651883 | 9.86e-02 |
| 48 | mutation_rate:Environmentsuc | 2.93e+05 | 1.84e+07 | 0.015939 | 9.87e-01 |

**Table S3:** Pairwise t-tests comparing the total number of mutations between strains, tested separately for each day. p-values were adjusted for multiple comparisons using the Benjamini–Hochberg false discovery rate procedure within each day separately (p_adj). p-values <0.05 are highlighted in bold.

| Day | Strain1 | Strain2 | t_stat | p_value | p_adj |
|---|---|---|---|---|---|
| 16 | ΔmutH | Δnth-nei | -0.14 | 8.96e-01 | 9.09e-01 |
| 16 | ΔmutH | ΔmutT | -1.01 | 3.58e-01 | 7.11e-01 |
| 16 | ΔmutH | WT | 8.71 | **3.31e-04** | **3.31e-03** |
| 16 | ΔmutH | ΔmutY | 0.12 | 9.09e-01 | 9.09e-01 |
| 16 | Δnth-nei | ΔmutT | -0.30 | 7.79e-01 | 9.09e-01 |
| 16 | Δnth-nei | WT | 3.00 | **3.02e-02** | 7.55e-02 |
| 16 | Δnth-nei | ΔmutY | 0.17 | 8.72e-01 | 9.09e-01 |
| 16 | ΔmutT | WT | 7.06 | **8.83e-04** | **4.41e-03** |
| 16 | ΔmutT | ΔmutY | 0.86 | 4.27e-01 | 7.11e-01 |
| 16 | WT | ΔmutY | -5.76 | **2.21e-03** | **7.38e-03** |
| 48 | ΔmutH | Δnth-nei | 2.40 | 6.13e-02 | 8.76e-02 |
| 48 | ΔmutH | ΔmutT | -7.13 | **8.44e-04** | **4.22e-03** |
| 48 | ΔmutH | WT | -1.49 | 1.98e-01 | 2.20e-01 |
| 48 | ΔmutH | ΔmutY | -2.28 | 7.17e-02 | 8.97e-02 |
| 48 | Δnth-nei | ΔmutT | -7.60 | **6.25e-04** | **4.22e-03** |
| 48 | Δnth-nei | WT | -2.63 | 4.65e-02 | 7.76e-02 |
| 48 | Δnth-nei | ΔmutY | -5.78 | **2.18e-03** | **7.26e-03** |
| 48 | ΔmutT | WT | 3.11 | **2.66e-02** | 5.33e-02 |
| 48 | ΔmutT | ΔmutY | 3.52 | **1.70e-02** | **4.25e-02** |
| 48 | WT | ΔmutY | -0.88 | 4.21e-01 | 4.21e-01 |

**Table S4:** Primers used for strain confirmation, 5’ to 3’.

| Gene | Forward primer (primer length) | Reverse primer (primer length) | Amplicon length |
|---|---|---|---|
| ***mutH*** | CAGAGAATTGAACAACGCATG (21) | GTAATTATATGCCCGGAAAGCG (22) | 818 |
| ***mutY*** | CAAGCATGATAAGGCCGTGG (20) | CTGCTTCACGTTGCAGGAAAGTAC (24) | 1219 |
| ***mutT*** | GCCTGCAATAAAAGCTAACTG (21) | TGAGTTAGAAATCCCTTTAGGC (22) | 555 |
| ***nth*** | TGCTGGCAGGAAAATACCTGATTGA (25) | TGATAACGCTAAGCAATGGCATCA (24) | 834 |
| ***nei*** | CGCCGACTACGCTCTGCATTT (21) | TGGCGATGCAAACCCTACGC (20) | 969 |
